# Autophagy activator AA-20 engages CDK5 to promote lipophagy and improve neurodegeneration-associated phenotypes

**DOI:** 10.64898/2026.10.30.760220

**Authors:** Ee Phie Tan, Lydia Ambaye, Anne-Laure Mahul-Mellier, Yuting Zhang, Patricia Martínez-Botía, Sergio R. Labra, Lynée A. Massey, Zi Gao, Zainab Ahmed, Karina Nugroho, Carl Ash, Wen Ren, Christian M. Cole, Yin Wu, Sarah Ly, Jadon Compher, Mireya Almaraz, Oren Lederberg, Alex Matthewson, Derek Rhoades, Caroline A. Cuoco, Danny Garza, Evan T. Powers, Joao A. Paulo, Daniel Finley, Steven P. Gygi, Michael Petrascheck, Stuart A. Lipton, Malene Hansen, Stefano Forli, Miguel A. Prado, Hilal A. Lashuel, Jeffery W. Kelly

## Abstract

Lipophagy, the selective degradation of lipid droplets (LDs) by autophagy, has the potential to ameliorate neurodegenerative diseases such as Parkinson’s (PD) and Alzheimer’s (AD), wherein LD accumulation has been linked to the exacerbation of the proteinopathy component of neurodegeneration. Pharmacological lipophagy inducers remain scarce. We previously identified AA-20 as a novel autophagy activator that acts independently of mTORC1 inhibition, thereby providing a means of stimulating autophagy while avoiding the immunosuppressive side effects associated with direct mTORC1 inhibition. Prior work also showed that AA-20 treatment reduces PolyQ aggregate levels in several *C. elegans* tissues. Here, we demonstrate that AA-20 enhances LD recruitment to autophagosomes and promotes LD degradation via lysosomal acid lipase activation in murine hippocampal neuronal cells, consistent with lipophagy activation. In human Swedish Amyloid Precursor Protein mutated cerebral organoids, two weeks of AA-20 treatment normalized aberrant lipidomic profiles by selectively reducing various disease-associated lipids, while also correcting the AD-associated electrical hyperexcitability phenotype. Moreover, we identified cyclin-dependent kinase 5 (CDK5), a neuronal kinase linked to AD hyperexcitability and PD pathology, as a target of AA-20. AA-20-induced autophagy and lipid clearance in HT22 cells require CDK5. In a neuronal seeding model of α-synuclein pathology, AA-20 reduced pS129-α-synuclein levels. Collectively, these findings indicate AA-20 can promote lipophagic clearance of LDs and mitigate multiple cellular phenotypes associated with lipidopathy and proteinopathy in neurodegenerative disease models.

**SIGNIFICANCE:** Neurodegenerative diseases such as Alzheimer’s and Parkinson’s are associated with impaired autophagy and abnormal lipid droplet (LD) accumulation. Few small molecules enhance neuronal lipid clearance without inhibiting mTORC1, but chronic blockade of which is associated with immunosuppressive and metabolic side effects. This study shows that AA-20, an mTORC1-independent autophagy activator, promotes lipophagic LD clearance in neuronal cells and reduces selected disease-associated lipids in human Alzheimer’s cerebral organoids in a lysosomal acid lipase-dependent manner. AA-20 binds CDK5 and appears to modulate, rather than classically inhibit, this neuron-enriched kinase. In an exploratory Parkinson’s neuron model, AA-20 also reduced pathological α-synuclein phosphorylation. These findings support further investigation of AA-20 as a tool to probe neuronal lipid clearance and neurodegeneration-associated phenotypes.

## INTRODUCTION

Macroautophagy, referred to here as ‘autophagy’, is a conserved process that sequesters cellular cargo into autophagic vesicles called autophagosomes (APs) that fuse with lysosomes to form autolysosomes (ALs), wherein lysosomal hydrolases degrade the contents and support cellular homeostasis (*1*). Among selective forms of autophagy, lipophagy refers to the selective targeting of lipid droplets (LDs) for lysosomal degradation. Lipophagy was initially discovered through the observation that autophagy inhibition causes accumulation of cellular triacylglycerol and cholesterol content as well as LDs, accompanied by co-localization of LD-containing APs with lysosomes (*2*). Although receptor-mediated LD targeting remains understudied (*3–6*), it is clear that lipophagy delivers LDs to autolysosomes for degradation by lysosomal acid lipase (LAL/LIPA) (**Fig. 1A**) (*7*).

**Fig. 1.**
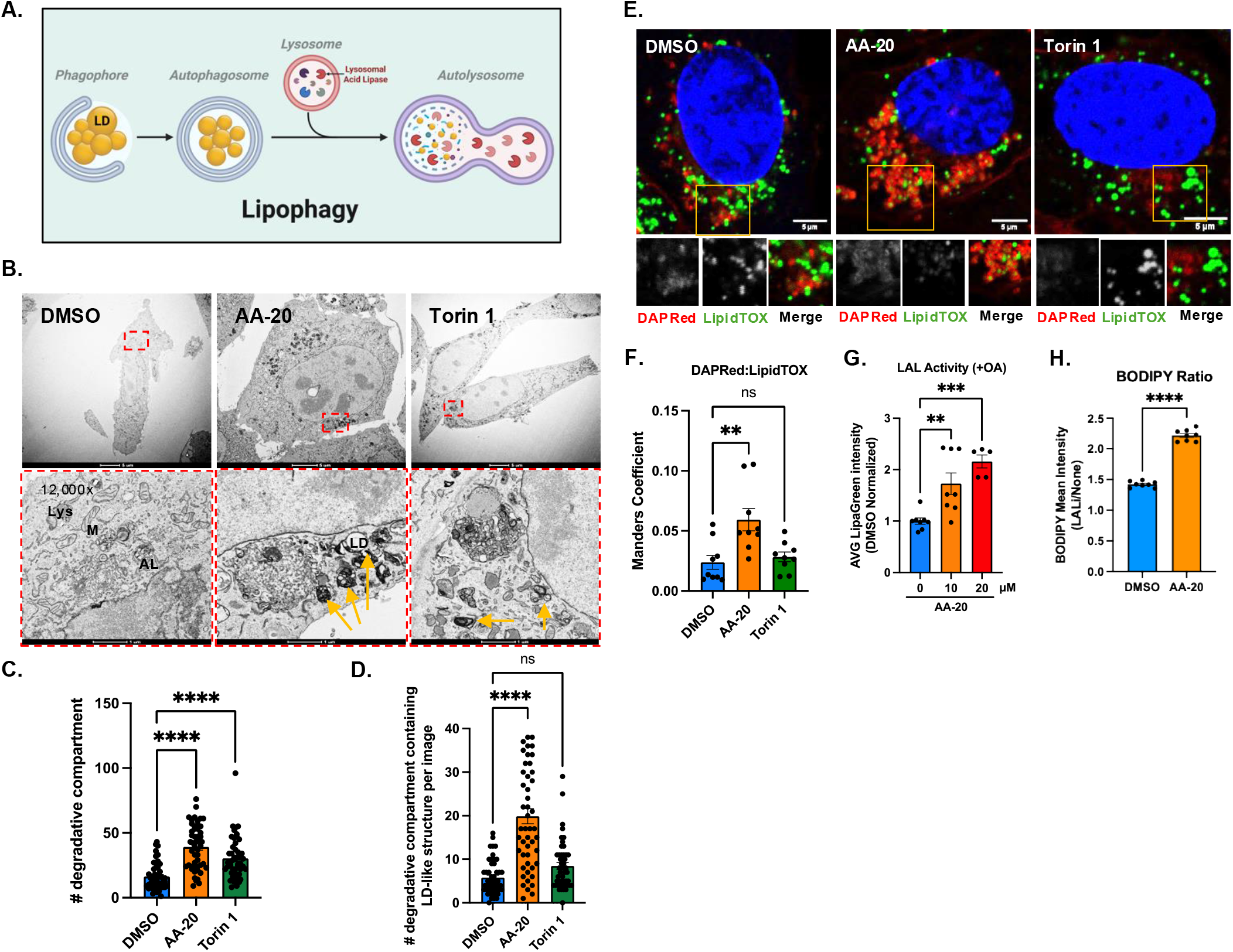
AA-20 enhances lipophagy in oleic acid-challenged HT22 cells. (A) Schematic depicting the process of lipophagy. Generated using BioRender. (B) Representative transmission electron micrographs of HT22 WT cells after ∼20 h pre-treatment with 60 μM Oleic Acid (OA)-BSA and 24 h treatment with DMSO (0.1%), 20 μM AA-20, and 150 nM Torin 1. (C) Quantification of (A), the number of electron-dense representing degradative compartments per cell (indicated by yellow arrows) and (D) the number of lipid droplet-containing degradative compartments per cell. *N=*50 cells per condition. \*\*\*\**p* < 0.0001, by ordinary one-way ANOVA. (E) Representative confocal micrographs of HT22 cells incubated with DAPRed (red), LipidTOX (green) and co-stained with Hoechst (blue) after 20 h pre-treatment of 60 μM OA-BSA and 24 h treatment with DMSO (0.1%), 150 nM Torin 1, or 20 μM AA-20. (F) Quantification of (E), DAPRed (autophagosome) and LipidTOX (LD) colocalization by Manders coefficient. *n=*3 independent experiments. Scale bar, 10 μm. (G) Quantification of average LipaGreen intensity after ∼20 h pre-treatment with 60 μM OA-BSA and 24 h DMSO (0.1%), 10 μM, or 20 μM AA-20 treatment in HT22 WT cells, normalized to DMSO. *n=*3 independent experiments, each with 2-3 biological replicates. \*\**p* < 0.01, \*\*\**p* < 0.001, by ordinary one-way ANOVA. (H) Neutral lipid ratio, calculated as the ratio of BODIPY fluorescence intensity from (SI Fig 1I), comparing conditions with and without the LAL inhibitor Lalistat 2, in the presence of either DMSO or AA-20 (DMSO+LALi/DMSO (blue bar), AA-20+LALi/AA-20 (orange bar). Error bars represent SEM. \*\*\*\**p* < 0.0001, by unpaired t-test.

Unc-51-like kinase 1 (ULK1) is a key kinase that initiates AP formation (*8, 9*). Under starvation conditions, mammalian target of rapamycin complex 1 (mTORC1) phosphorylates ULK1 at Ser757 and thereby suppresses the ULK1 complex, preventing autophagy initiation (*10*). Because mTORC1 also regulates mRNA translation and immune function (*11*), prolonged pharmacological inhibition (e.g., by rapamycin, an FDA-approved immunosuppressant with anti-aging effects) can lead to side effects such as immunosuppression and glucose intolerance (*12*). mTORC1-inhibition-independent autophagy enhancers therefore represent alternative strategies for treating age-related neurodegeneration (*13*). To identify such compounds, we previously screened ∼1 million small molecules for those that reduced the number of LDs in human SW1990 adenocarcinoma cells and through secondary screens identified 24 autophagy activators that accelerate LD clearance without inhibiting mTORC1, including AA-20 (*14, 15*). AA-20 extends lifespan in *C. elegans* and clears both LDs and protein aggregates in an autophagy-dependent manner in *C. elegans* and human cells (*15*). However, the mechanism of AA-20 remains unknown, and it is unclear whether its ability to accelerate LD clearance reflects true induction of lipophagy.

Aberrant LD accumulation and reduced lipophagic clearance can occur with aging (*16*), and experimentally impairing lipophagy can promote neurodegeneration (*17, 18*). In Parkinson’s disease (PD) models, neurons accumulate LDs (*19–22*). α-synuclein (aSyn), a lipid-binding protein that aggregates into Lewy bodies (LBs), can be degraded in part by autophagy (*23*), and electron microscopy of PD donor tissue shows that aSyn-immunopositive LBs and neurites contain abundant lipids together with lysosome- and AP-like structures (*24*). These findings suggest a possible relationship between lipid dysregulation and aSyn pathology, although whether enhancing lipophagy can influence aSyn-related toxicity remains largely untested. Pharmacological agents capable of promoting lipophagy in neurons are scarce (*25*), limiting direct experimental tests of this idea.

Cyclin-dependent kinase 5 (CDK5) is an atypical CDK activated by p35 or p39 rather than cyclins. It regulates neuronal development, synaptic plasticity, learning, memory, and excitability (*26*). CDK5 dysregulation is implicated in AD, PD, and Huntington’s disease (*27–30*). Its role in autophagy regulation appears context-dependent. Bioinformatic analysis predicts that CDK5 phosphorylates ULK1 (*31*). In *Drosophila melanogaster*, CDK5 has been proposed to support basal autophagy in an Atg1/ULK1-dependent manner and to be protective in early neurodegenerative stress (*32, 33*). In contrast, in human cells, CDK5 activated by p25 (a pathologic truncated form of p35), can inhibit autophagy by phosphorylating Vps34, a lipid kinase required for autophagosome formation (*34*). These opposing findings underscore the need for further mechanistic and therapeutic investigation of CDK5 in autophagy.

In this study, we show that the autophagy activator AA-20 promotes lipophagy in a largely CDK5-dependent manner, although additional targets cannot be excluded, distinguishing it from canonical mTORC1 inhibitors. In human hereditary AD cerebral organoids, multidose AA-20 treatment attenuates electrical hyperexcitability and restores both the phosphoproteome and lipidome toward wild-type-like states, with lipidome normalization appearing to depend on LAL/LIPA. In a neuronal model of PD (*35*), where aSyn aggregation is linked to interactions with lipids and membranous organelles (*24, 36, 37*), treatment with AA-20 results in significant reduction of the levels of pS129-aSyn pathological aggregates. Collectively, these findings highlight three advances, AA-20 can: (1) promote lipophagic clearance in neuronal-like cells; (2) correct disease-associated lipid abnormalities in an AD organoid model; and (3) reduce pS129-aSyn levels in a PD neuronal model. Prior studies have linked LD accumulation and lipophagy to aSyn pathology (*20, 38*), although the present study did not determine whether the reduction in pS129-aSyn was mediated by lipophagy. These results support further investigation of AA-20 as a compound that can improve selected features of lipid dysregulation and aSyn pathology in neuronal models.

## RESULTS

### Autophagy activator AA-20 promotes lipophagy in neuronal-like HT22 cells

AA-20 (**SI Fig. 1A**) is a small molecule autophagy activator (*15*), previously discovered through a high-throughput screen (HTS) to identify compounds that increase cellular lipid droplet (LD) clearance (*14*) (**SI Fig. 1A**). A 1,3,5-trisubstituted triazine comprises the core (colored black) of AA-20, with two 4-methoxyaniline substituents (colored blue) and one benzothiazole substituent (colored red) each attached to the triazine sp^2^-hybridized carbons (**SI Fig. 1A**) (*15*). Building on our prior findings that AA-20 induces autophagy in both HeLa cells and *C. elegans* (*15*), we sought to further scrutinize its effects in a neuronal context. To this end, we used the immortalized HT22 cell line, which retains key features of differentiated hippocampal neurons and is widely used to model neuronal autophagy and stress responses relevant to neurodegeneration (*39*).

To evaluate autophagy changes, we used HT22 cells stably expressing the GFP-LC3B-RFP-LC3BΔG ratiometric probe (*40*). Briefly, ATG4 cleavage enables GFP-LC3B lipidation and phagophore/AP association; GFP quenches upon lysosomal fusion to autolysosome (AL), while RFP-LC3BΔG, cleavage product lacking the C-terminal glycine, cannot be lipidated and remains cytoplasmic as an internal RFP control. A decreased GFP/RFP fluorescence ratio measured in a flow cytometry experiment indicates autophagy activation (*40*). AA-20 treatment (1–20 µM) dose-dependently reduced the GFP/RFP ratio compared to vehicle (DMSO) over 24 h, with significant effects observed at concentrations >10 µM (**SI Fig. 1B**). We also measured lysosomal function by assaying cathepsin D (CTSD) activity 24 h after AA-20 (10 μM) treatment commencement. Like the known autophagy activator Torin 1, AA-20 significantly increased CTSD activity, while Bafilomycin A1 (BafA), a lysosomal acidification inhibitor, reduced it (**SI Fig. 1C**). This finding aligns with our prior evidence showing conserved autophagy induction by AA-20 (*15*).

Lipophagy recruits LDs into APs for degradation in ALs (*2*). We previously showed that AA-20 promotes autophagy-dependent LD clearance in non-neuronal human cells and multiple *C. elegans* tissues (*15*). Here, we sought to investigate whether AA-20 induces lipophagy in the neuronal HT22 cell model. Cells were pre-treated with 60 µM oleic acid (OA) for 16h to increase LD content (*15*), then treated with AA-20 and the mTORC1 inhibitor Torin 1. BODIPY labeling of LDs, as in our prior HTS study (*14*) showed that AA-20 significantly reduced total LD area (**SI Fig. 1D-E**). Negative stain transmission electron microscopy (TEM) was also used to examine degradative compartments. ALs and lysosomes appear as electron-dense structures with amorphous content due to their acidic interior (*41–43*). Manual quantification of these electron-dense compartments (**Fig. 1B**, marked with yellow arrows) showed that both AA-20 and Torin 1 significantly increased their number (**Fig. 1B-C**), consistent with enhanced autophagy. LD containing degradative compartments are depicted as electron deficient (white) circles. Notably, AA-20-treated cells exhibited a significant increase in LD-containing degradative compartments.

In contrast, Torin 1-treated cells did not in the 50 cells examined (**Fig. 1D**), despite their similar induction of degradative AL-like compartments overall (**Fig. 1C**). Consistent with this finding, fluorescence imaging with the AP marker DAPRed (a commercial fluorescent dye used to label APs (*44*)) and the LD marker LipidTOX showed that AA-20, but not Torin 1, increased LD-AP colocalization in HT22 cells (**Fig. 1E-F**). Together, these results indicate that AA-20 promotes LD recruitment into APs/ALs in OA-pretreated neuronal HT22 cells, supporting its ability to enhance lipophagy.

LAL/LIPA is the primary enzyme that hydrolyzes cholesterol esters and triglycerides during lipophagy (*45, 46*). Using a LysoLive assay in which the enzyme substrate LipaGreen fluoresces upon LAL/LIPA-mediated cleavage in live cells, we found that AA-20 significantly increased LAL activity by ∼1.5-fold at 10 µM and ∼2-fold at 20 µM treatment, respectively) in both OA-BSA-pretreated cells (**Fig. 1G**) and untreated HT22 cells (**SI Fig. 1F**), consistent with our prior observation of increased LysoTracker Red intensity in HeLa cells treated with AA-20 (*15*). The increase in HT22 LAL activity occurred without changes in LAL protein levels (**SI Fig. 1G-H**), indicating elevated specific activity. This enhancement of LAL activity in OA-untreated cells suggests that AA-20 may prime lipophagy pathways for basal lipid turnover, while its effect in OA-pretreated cells points to a role in promoting LD degradation under lipid stress.

Our prior findings in *C. elegans* and SW1990 cells demonstrated that AA-20-mediated LD clearance is, at least in part, mediated, via autophagy (*15*). To test whether AA-20-mediated LD clearance requires LAL/LIPA activity, we employed Lalistat 2 inhibition (LALi). AA-20 alone significantly reduced LD levels compared to control, but this reduction was blocked by inhibitor LALi (**SI Fig. 1I**). We next quantified LAL-dependent LD turnover in two ways: the ratio of LD levels with versus without LALi (DMSO+LALi / DMSO (blue bar), AA-20+LALi / AA-20 (orange bar)), **Fig. 1H**) and the differences in LD levels between LALi-treated and untreated conditions (DMSO+LALi – DMSO (blue bar), AA-20+LALi – AA-20 (orange bar)) **SI Fig. 1J**). Both measures showed greater LAL dependence in AA-20 treated cells than in controls (**Fig. 1H, SI Fig. 1J**), indicating that AA-20 does promote LD clearance in an LAL-dependent manner.

### AA-20 normalizes lipid dysregulation and hyperexcitability AD cerebrocortical organoids

Human AD brains exhibit excessive LDs enriched in triacylglycerols (TGs), diacylglycerols (DGs), cholesteryl esters (Chol-esters), and ceramides (*47–49*). We therefore tested the effects of AA-20 treatment in human induced pluripotent stem cell (iPSC)-derived cerebrocortical organoids carrying the AD-associated heterozygous Swedish Amyloid Precursor Protein mutation, comparing to isogenic wild-type (WT/WT) organoids as controls (*50*) (**Fig. 2A**). These organoids formed organized cortical layers (**SI Fig. 2A**) and accumulated fibrillar phosphorylated-tau (pTau, AT8) by 3 months, which was absent in WT controls (**SI Fig. 2B**), consistent with previous reports that this model recapitulates key features of cortical development and early AD pathology (*51, 52*). Multielectrode array (MEA) recordings showed that AD organoids developed hyperexcitability over time, also recapitulating early AD pathology (*52–55*). AA-20 reduced this hyperexcitability phenotype in AD organoids without significantly affecting WT organoids (**Fig. 2B**). Heterozygous APPswe/WT organoids also had lower electrode impedance (viability index) compared to WT organoids (**SI Fig. 2C, cf. DMSO conditions, blue**). AA-20 rescued this impedance reduction, indicating a broader protective effect.

**Fig. 2.**
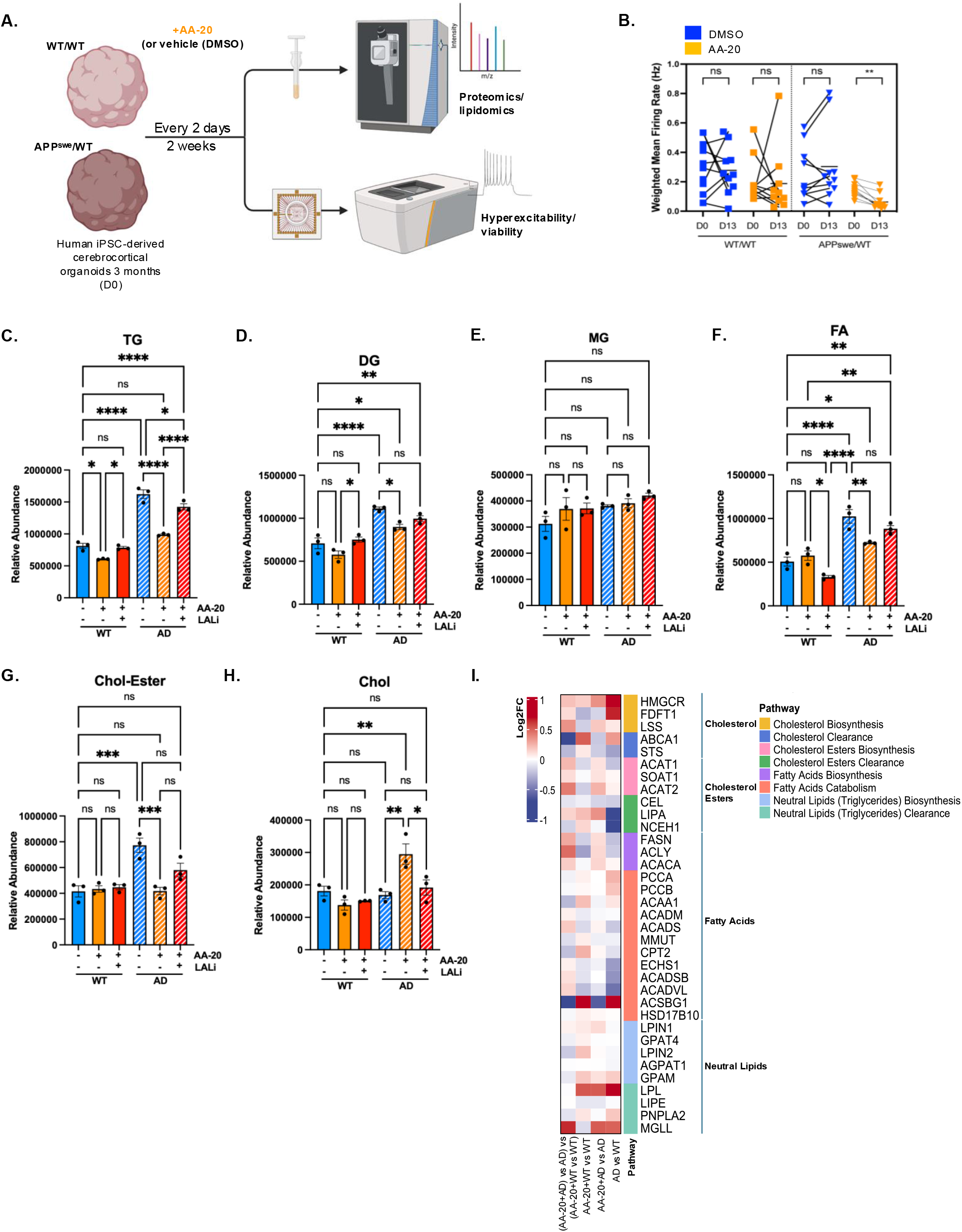
Chronic AA-20 treatment normalizes AD-associated pathology in human iPSC-derived cerebral organoids. (A) Schematic illustrating experimental setup and treatment timeline for human cerebral organoids comparing AD model (APPswe/WT) vs. wild-type control (WT/WT). Created using BioRender. (B) Quantification comparing the organoid neuronal activity before (D0) and after the 13-day treatment (D13), as calculated by the weighted mean firing rate (Hz) in n = at least 10 organoids per condition. (C-H) Plots depicting the relative abundance of selected lipid species from lipidomic analysis after two weeks of treatment with 5 μM AA-20 or co-treatment with 5 μM Lalistat 2, LAL inhibitor, in WT or AD organoids. Lipid species include triacylglycerols (TG) (C), diacylglycerols (DG) (D), monoglycerols (MG) (E), fatty acids (FA) (F), cholesterol ester (Chol-Ester) (G), and cholesterol (Chol) (H). *n*=3 biological replicates. All error bars represent SEM. Statistical significance: \**p* < 0.05, \*\**p* < 0.01, \*\*\**p* < 0.001, \*\*\*\**p* < 0.0001, determined by two-way ANOVA with Tukey’s multiple comparisons test. Also see Supplemental Data 1 for the complete dataset. (I) Heatmap depicting the log_2_-fold changes of those enzymes related to lipid metabolism that were detected by TMT proteomics. Enzymes were ordered and colored according to the specific lipid pathway they belonged to. Conditions are as follows: “AD”, APPswe/WT; “WT AA-20”, WT AA-20/WT; “AD AA-20”, APPswe AA-20/APPswe; and “AD AA-20 – WT AA-20”, APPswe AA-20/APPswe vs. WT AA-20/WT. Refer to Supplemental Data 3-4 for the completed dataset.

To profile the proteomic and lipidomic effects of AA-20, we performed quantitative unenriched TMT-based proteomics and lipidomics on 3-month-old WT and APPswe/WT organoids treated every other day for 14 days with AA-20 (final concentration of 5 µM (0.1% DMSO) (**Fig. 2A**). Lipidomics showed that AD organoids had elevated TGs, DGs, fatty acids (FAs), and Chol-esters levels compared to gene corrected isogenic WT organoids (**Fig. 2C–G**, **Supplemental Data 1**). In WT organoids, AA-20 selectively reduced TG (key lipophagy substrate), and this effect was blocked by the LAL inhibitor (LALi) (**Fig. 2D**). In AD organoids, AA-20 significantly reduced TGs, DGs, FAs, and Chol-esters, but did not affect monoglycerols (MGs) (**Fig. 2D–G**). These reductions were substantially attenuated by LALi (**Fig. 2C–G**). AA-20 also increased free cholesterol levels in AD organoids (**Fig. 2H**) in a LAL-dependent manner, consistent with enhanced LD Chol-ester hydrolysis (*56*) (**Fig. 2G**). Thus, AA-20 produced an LAL-dependent correction of lipid dysregulation in the AD background that is smaller in WT organoids.

To further elucidate the likely mechanisms driving the observed lipid alterations, we analyzed the whole proteome (refer to **Supplemental Data 2** for TMT information). We used statistical comparisons to distinguish disease effects (AD vs WT), direct AA-20 effects in AD (AA-20+AD vs AD), AA-20 effects in WT (AA-20+WT vs WT), and disease-selective drug responses via the interaction term ((AA-20+AD vs AD) – (AA-20+WT vs WT)) (**Supplemental Data 2–4**). Untreated AD organoids showed reduced expression of lipid clearance enzymes (CEL, LIPA/LAL, NCEH1). AA-20 upregulated these enzymes to a greater extent in AD than in WT organoids, with the interaction contrast confirming disease-preferential induction, most notably of LIPA/LAL (**Fig. 2I**, **Supplemental Data 3-4**).

Pathway-level analysis was consistent with these lipid changes. In cholesterol metabolism, untreated AD organoids showed increased biosynthetic enzymes and reduced clearance enzymes, yet steady-state free cholesterol was maintained (**Fig. 2I**, **Supplemental Data 3-4**). In the AD background, AA-20 left biosynthesis largely unchanged while further reducing clearance-enzyme expression (**Fig. 2I**, **Supplemental Data 3-4**), consistent with the observed increase in free cholesterol (see **Fig. 2H**). In FA metabolism, catabolic enzymes were downregulated in untreated AD organoids; AA-20 restored these enzymes in a disease-preferential manner, while biosynthetic enzymes showed only a modest upward trend (**Fig. 2I**, **Supplemental Data 3–4**). Together, the lipidomics data demonstrate LAL-dependent correction of lipid dysregulation by AA-20, while the proteomics data indicate that this effect is accompanied by disease-selective upregulation of lipid clearance pathways. Collectively, having observed disease-selective effects of AA-20 in AD organoids, we returned to HT22 cells to compare its mechanism with that of a canonical mTORC1 inhibitor.

### AA-20 alters HT22 cell proteomic profiles differently from an mTORC1 inhibitor

We previously found that AA-20 appears to induce autophagy without inhibiting mTORC1 (*15*). To further investigate this mechanism, we performed tandem-mass-tagging (TMT) proteomic (whole-proteome) and phosphoproteomic analyses in HT22 cells treated with either AA-20 or Torin 1 for 24 h (refer to **Supplemental Data 2** for TMT information). Whole-proteome analysis quantified 7,529 proteins. AA-20 altered 1,898 proteins (921 proteins up-regulated, 977 proteins down-regulated), whereas Torin 1 altered 3,117 proteins (1,638 up-regulated, 1,479 down-regulated) (FDR < 0.05; **SI Fig. 3A**, **Supplemental Data 5**). Phosphoproteomic analysis quantified >5,000 phosphosites, of which 796 sites were altered by AA-20 (398 up-regulated, 398 down-regulated) versus 1,525 phosphosites altered by Torin 1 (804 up-regulated, 721 down-regulated) (FDR < 0.05; **SI Fig. 3B**, **Supplemental Data 6**). Thus, Torin 1 produced a broader proteomic and phosphoproteomic response than AA-20.

**Fig. 3.**
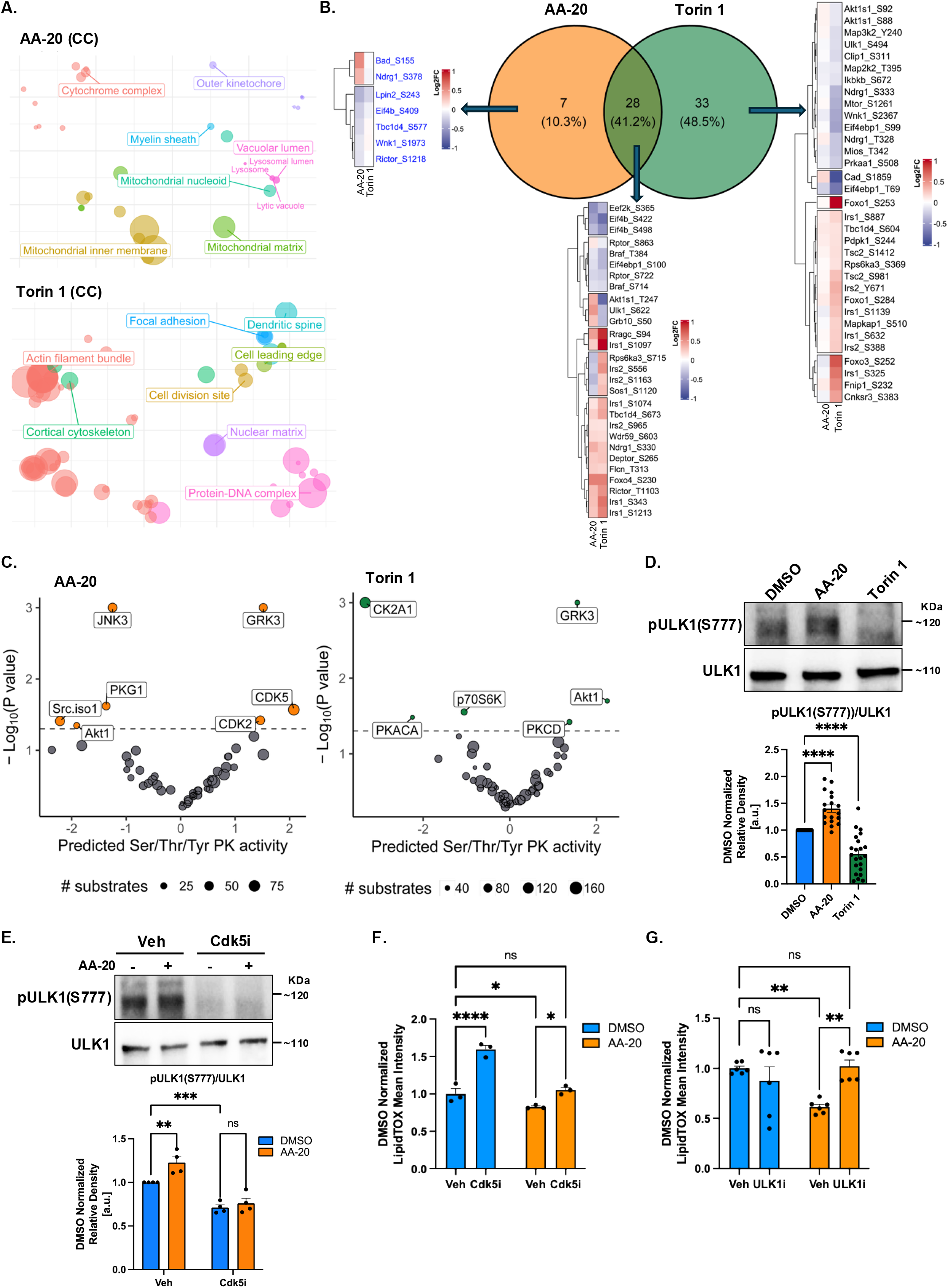
AA-20 distinctly modifies HT22 cell proteomic profiles compared to an mTORC1 inhibitor and promotes ULK1 phosphorylation via CDK5-dependent mechanisms. (A) Principal component analysis (PCA) of gene ontology (GO) terms (cellular compartment, CC) that are significantly enriched in the AA-20 (20 µM) and Torin 1 (150 nM)-treated HT22 cells, compared to DMSO treatment alone. Only significantly up-regulated proteins were used. To reduce redundancy, grouping of terms was performed by semantic similarity among GO terms. Boxed labels and accompanying colors represent the resulting broader or parent terms. The size of the points represents the similarity scores. Refer to Supplemental Data 7 for the complete list of the GO terms. (B) Schematic displaying the overlapping mTOR pathway phosphorylation sites significantly altered as a result of AA-20 or Torin 1 treatment, analyzed from both phosphoproteomic data and known *mus musculus* mTOR pathway proteins. Hierarchical clustering of the heatmaps relied on the Euclidean distance and the complete method. (C) Prediction of kinase activity from phosphoproteomic data using KinSwing package in R. Predicted kinases significantly altered by AA-20 or Torin 1 treatment are displayed as a volcano plot of reduced to increased kinase activity as a function of statistical significance (-Log_10_(P value)) (refer to Supplemental Data 9 for complete dataset). All proteomic analyses were performed with four biological replicates. (D-E) Representative immunoblots and densitometry analyses of pULK1-S777, normalized to total ULK1 protein level and DMSO control. (D) HT22 cells were treated with DMSO, AA-20 (10 µM), or Torin 1 (150 nM) for 24 h respectively. *n*=18 independent replicates. \*\*\*\**p*<0.0001, by ordinary one-way ANOVA. (E) µM AA-20 and co-treated with either vehicle (DMSO) or 5 µM 20-223, a CDK5 inhibitor, for 24 h. *n*=4 independent replicates. **p<0.01, ***p<0.001, by two-way ANOVA with Tukey’s multiple comparisons. (F-G) Bar graph showing the mean intensity of LipidTOX by flow cytometry in HT22 cells. Cells were preincubated with 60 μM OA-BSA, then treated with either DMSO (0.2%), (F) 5 μM 20-223 (*n*=3 biological replicates), or with (G) 5 μM of ULK-101 alone (*n*=6 biological replicates), or co-treated with 10 μM AA-20, respectively. Statistical significance: ns, not significant; \*\**p* < 0.01, \*\*\**p* < 0.001, \*\*\*\**p* < 0.0001, by ordinary one-way ANOVA. All error bars represent SEM.

Gene Ontology (GO) cellular components (CC) enrichment showed that AA-20 uniquely upregulated mitochondrial terms (e.g., cytochrome complex, mitochondrial nucleoid, mitochondrial inner membrane and matrix), and vacuolar lumen (e.g., lysosomal lumen, lysosome and lytic vacuole), a profile distinct from Torin 1 (**Fig. 3A, Supplemental Data 7**). This lysosomal enrichment (see **Fig. 3A**) is consistent with our prior finding that AA-20 increases lysosomal abundance and function in HeLa cells (*15*).

To test whether AA-20 modulates the mTOR pathway, we examined phosphosites on 68 curated *Mus musculus* mTOR pathway substrate proteins drawn from a literature list of 191 (*57–60*) (refer to **Supplemental Data 8** for the full list). AA-20 altered 7 unique phosphosites, compared to Torin 1 with 33 unique phosphosites and 28 shared by both AA-20 and Torin 1 (**Fig. 3B**). Some shared changes were compatible with reduced mTOR signaling, but many were not mTORC1-specific or directly linked to autophagy. Immunoblotting (IB) confirmed that, unlike Torin 1, AA-20 did not change phosphorylation of the canonical mTORC1 substrates pS757-ULK1 or pT389-S6K (**SI Fig. 3C**) in HT22 cells, consistent with prior HeLa cell findings (*15*).

Unbiased KinSwing analysis (see Methods) of the phosphoproteome (**Supplemental Data 6**, **9**) inferred increased CDK2 and CDK5 activity after AA-20 treatment, together with reduced c-Jun N-terminal kinase 3 (JNK3)/Mitogen-Activated Protein Kinase 10 (MAPK10) and Protein Kinase G1 (PKG) activity (**Fig. 3C**, **Supplemental Data 9**). Torin 1 instead reduced S6 kinase 1 (S6K1) activity, and increased Akt1 activity (*61*) (**Fig. 3C**). Both compounds were associated with inferred G-protein-coupled receptor kinase 3 (GRK3) activity (**Fig. 3C**). Thus, AA-20 and Torin 1 engage distinct kinase programs, consistent with an mTORC1-independent mechanism for AA-20.

### AA-20 enhances ULK1 phosphorylation and lipophagy via CDK5/2-dependent mechanisms

CDK5 is the predominant CDK in post-mitotic neurons and regulates neuronal development and synaptic plasticity (*26*), whereas CDK2 mainly drives cell cycle progression (*62*). Given their reported roles in autophagy (*63*), we examined CDK2- and CDK5-associated phosphosites in the HT22 phosphoproteome (**Supplemental Data 7**). Of 61 identified phosphosites, 12 (19.7%) were assigned to CDK2, 23 (37.7%) to CDK5, and 26 (42.6%) were shared. The shared sites included ULK1 Ser622 (equivalent to human ULK1 Ser623; UniProt O75385/O70405) (**SI Fig. 3D**, **Supplemental Data 10–11**). Although it remains unclear whether CDK5 directly phosphorylates ULK1/Atg1, CDK5 can influence ULK1/Atg1 phosphorylation and autophagy in *Drosophila* (*33*). AA-20, but not Torin 1, increased pS622-ULK1 levels according to IBs (**SI Fig. 3E**). We also examined pS777-ULK1, an AMPK-regulated autophagy activation site that can operate independently of mTORC1 inhibition under glucose-rich conditions (*10*). This site was not detected by phosphoproteomics, but IB showed that AA-20 significantly increased pS777-ULK1, whereas Torin 1 reduced it (**Fig. 3D**). The CDK5/2 inhibitor 20-223 (5 µM) reduced basal pS777-ULK1 and produced a non-significant 14.0% decrease in pS622-ULK1 (**Fig. 3E, SI Fig. 3F**). Co-treatment with 20-223 blocked the AA-20-induced increases in both pS622- and pS777-ULK1 levels (**Fig. 3E**, **SI Fig. 3F**). These findings indicate that AA-20 promotes ULK1 phosphorylation at Ser777 and Ser622 in a CDK5/2-dependent manner, consistent with the KinSwing inference of increased CDK5/CDK2 activity (refer to **Fig. 3C**, **SI Fig. 3E**).

To test whether this pathway is required for AA-20-induced autophagy, we used HT22 cells expressing the GFP-LC3B-RFP-LC3BΔG fluorescence autophagy reporter (see also **SI Fig. 1B**). AA-20 dose-dependently decreased the GFP/RFP ratio indicating increased autophagic flux, and this effect was blocked by 20-223 (5 µM) (**SI Fig. 3G**). AA-20 also increased CTSD activity (**SI Fig. 3H**, see also **SI Fig. 1D**). Since 20-223 alone substantially reduced basal CTSD activity, the influence of AA-20 in the presence of the CDK5/2 inhibitor 20-223 was difficult to interpret (**SI Fig. 3H**). Because ULK1 influences AL function in mammalian cells (*64*), we next used the ULK1 inhibitor ULK-101 (5 µM). ULK-101 alone did not alter CTSD activity but prevented AA-20 from increasing it (**SI Fig. 3H**). Thus, AA-20-induced autophagy appears to require CDK5/2-dependent ULK1 activity (refer to **Fig. 3D-E**, **SI Fig. 3E-H**).

We next asked whether the same pathway is required for AA-20-mediated LD reduction (**Fig. 2**). The CDK5/2 inhibitor 20-223 increased basal LD levels and prevented AA-20 from fully reducing LDs (**Fig. 3F**). ULK-101 likewise blocked AA-20-induced LD reduction (**Fig. 3G**). These results indicate that CDK5/2 contributes to both basal LD turnover and the AA-20 response, whereas ULK1 is specifically required for AA-20-mediated LD clearance. Collectively, AA-20 promotes autophagy and lipophagy in HT22 cells, at least in part through CDK5/2-dependent ULK1 activation, rather than through mTORC1 inhibition.

### AA-20 binds to CDK5 directly

Because our phosphoproteomics inferred increased CDK5 and CDK2 activity after AA-20 treatment (see **Fig. 3C**), we hypothesized that AA-20 may bind one or both kinases. We prioritized CDK5 as it is highly expressed in neurons, including in the hippocampus, and has an established role in neuronal autophagy (*65*), whereas CDK2 is more closely associated with cell cycle progression in proliferating cells. CDK5 and CDK2 share a conserved kinase fold and similar ATP-binding clefts (*66, 67*), so some ligand overlap is possible. We therefore asked whether AA-20 binds CDK5 directly.

A NanoBRET kinase binding assay (conducted by Promega) in live and lysed HEK293 cells expressing NanoLuc-fused CDK5 with the physiological activator p35 (CDK5r1) showed that AA-20 dose-dependently displaced the CDK5-specific tracer (IC_50_: 1.53 μM in live cells; 9.25 μM in lysed cells (**Fig. 4A-B**)). These cellular IC_50_ values are higher than the biochemical affinity, as is tyical for competitive displacement assays performed in the presence of ATP and cellular complnents. Cellular thermal shift assays in HT22 lysates (our main neuronal model) showed that AA-20 increased the CDK5 melting temperature from 38.9°C to 46.5°C (**SI Fig. 4A-B**), consistent with binding-induced stabilization. Surface Plasmon Resonance assays (*68*) with immobilized human recombinant CDK5/p35 further demonstrated high-affinity binding (Kd = 80 ± 31 nM) (**Fig. 4C**). These orthogonal assays indicate that AA-20 binds CDK5/p35 directly, providing a plausible mechanism for the observed CDK5-associated ULK1 phosphorylation (see **Fig. 3D-E**, **SI Fig. 3E-F**).

**Fig. 4.**
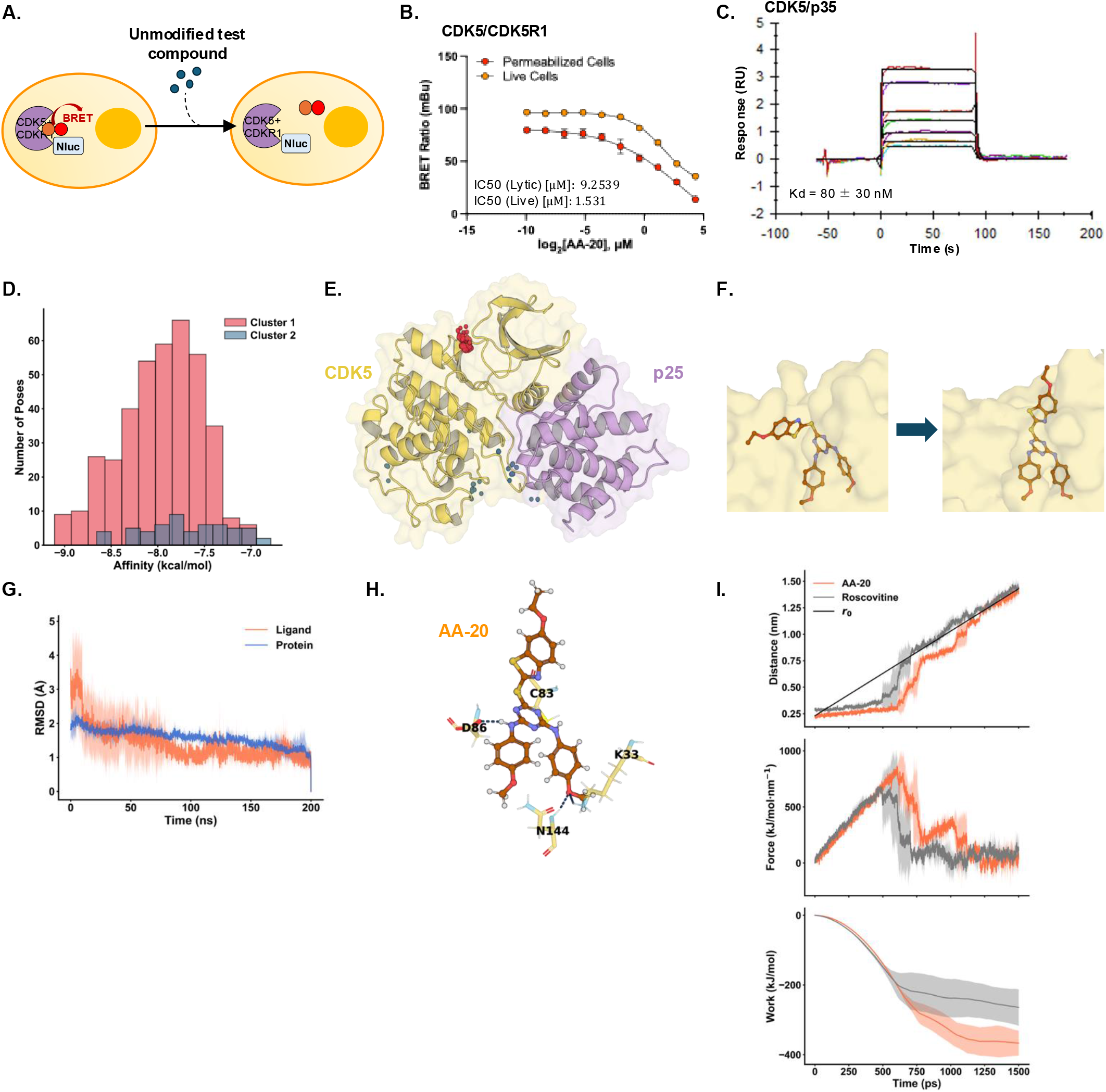
AA-20 acts as a CDK5 modulator. (A) Schematic of the Live-Cell Kinase NanoBRET Target Engagement Assay conducted by Promega. A cell-permeable energy transfer tracer reversibly binds to the kinase’s active site, producing a BRET signal in live cells. Binding of an unlabeled test compound to the selected kinase leads to a reduction in the BRET signal. (B) BRET ratios (mBu) were measured using AA-20 at concentrations ranging from 0 to 20 μM in HEK293T cells transiently expressing CDK5+CDK5R1 NLuc fusion proteins. Experiments were conducted in either lysed or live cells. (C) Representative SPR sensorgram showing SPR analysis of AA-20 binding to the CDK5/p35 complex. Response units (RU) on the y-axis versus time (s) on the x-axis, depicting the association and dissociation phases of AA-20 binding. The equilibrium dissociation constant (Kd) was determined to be 79 ± 31 nM, calculated from the kinetic parameters of the binding curves. *n*=3 independent repeats. (D) Distribution of predicted binding affinities for all 450 docking poses of AA-20 to the CDK5/p25 complex. Binding affinity (kcal/mol) is plotted on the x-axis, and the number of poses per affinity bin on the y-axis. Poses are colored by cluster: red for the ATP-binding pocket (Cluster 1) and blue for the region near the activation loop (Cluster 2). (E) Mapping of the two major pose clusters onto the CDK5/p25 complex structure. Red and blue dots represent the centers of mass of individual AA-20 poses in Cluster 1 and Cluster 2, respectively. (F) Binding pose of AA-20 in the CDK5/p25 complex before and after MD refinement. Left: top-ranked docking pose based on the highest affinity from blind docking. Right: representative bound conformation sampled from MD trajectories. (G) RMSD of AA-20 (orange) and CDK5/p25 backbone (blue) over time during MD simulations. Time (ns) is on the x-axis, RMSD (Å) on the y-axis. Lines show the mean across three replicas; shaded areas indicate ± standard deviation. (H) Representative MD-refined binding pose of AA-20 in the CDK5/p25 ATP-binding pocket. Hydrogen bonds between AA-20 and surrounding residues are shown as dashed lines. (I) Steered MD results comparing the unbinding of AA-20 and Roscovitine from the CDK5/p25 ATP-binding pocket. Top to bottom: ligand–protein center-of-mass distance, applied force, and cumulative work, all plotted over time. Lines show the mean across three replicas; shaded areas indicate standard deviation. r0 indicates the time-dependent target position of the harmonic restraint.

### Docking simulations indicate AA-20 may bind the CDK5 complex at its ATP-binding pocket

Given the evidence for direct CDK5 binding, we next used blind docking and 200 ns molecular dynamics (MD) simulations to explore and predict how AA-20 might engage the CDK5 complex. Simulations were based on the CDK5/p25 crystal structure (PDB ID: 1UNL) (*69*). p25 is a stable fragment of the physiological activator p35; the structure also contains a D144N mutation used to facilitate crystallization (*69*). Our AutoDock Vina docking (*70*) generated 450 poses with predicted affinities of –9.11 to –6.79 kcal/mol (**Fig. 4D**). Clustering showed that two primary clusters emerged (**Fig. 4E**, red and blue). The majority (394 poses, 88%) localized to the ATP-binding pocket, the canonical site used by many kinase ligands, whereas 56 poses (12%) mapped near the activation loop (**Fig. 4E**).

To evaluate the binding stability of AA-20 in the CDK5/p25 complex, we performed time-resolved molecular dynamics (MD) simulations after the static docking analysis. MD simulations starting from the best-scoring ATP-site pose showed that AA-20 remained in the pocket, with the benzothiazole group shifting toward the solvent-exposed entrance and then stabilizing. This rearrangement coincided with stabilization of AA-20 in the pocket by 100 ns (low root-mean square-deviation, RMSD) and limited ligand flexibility (root-mean-square fluctuation (RMSF) <1.5 Å, mainly at the benzothiazole) (**Fig. 4F-G**, **SI Fig. 4C-D**). Persistent hydrogen bonds were observed with Lys33, Asp86, and Asn144 (**Fig. 4H**). Lys33 (*71–73*) and Asp86 (*69, 74–78*) are conserved residues commonly contacted by CDK5 inhibitors. The Asn144 contact depends on the crystallization mutant and may not be present in WT CDK5.

Because this pose overlaps with the ATP site used by CDK5 inhibitor Roscovitine (*79*), we compared their unbinding resistance by steered MD. AA-20 (from MD endpoint) and Roscovitine (crystallographic pose, PDB ID: 1UNL) showed similar profiles, with AA-20 displaying slightly higher peak forces and work (**Fig. 4I**, **SI Fig. 4E**). These differences are within typical steered-MD variability and suggest that AA-20 can occupy the pocket as a classical inhibitor.

Since our MD simulations predict that AA-20 can stably occupy the CDK5 ATP-binding pocket, akin to Roscovitine, we next asked whether this binding inhibits CDK5. In ADP-Glo assays, which quantify ADP generated during phosphorylation of histone H1 by CDK5/p35 (Promega), Roscovitine potently inhibited activity, whereas AA-20 caused only partial inhibition near 40 μM (**SI Fig. 4F**). Thus, despite high-affinity binding to the CDK5 complex (refer to **Fig. 4B-D**), AA-20 did not behave as a classical CDK5 inhibitor. A more likely interpretation is that it modulates CDK5 activity or substrate engagement (see model in **Fig. 5**). This is consistent with AA-20 increasing CDK5/2-dependent ULK1 phosphorylation and with a CDK5/2 inhibitor blocking AA-20-induced autophagy and LD clearance in HT22 cells (see **Fig. 3D-G**, **SI Fig. 3E-H**).

**Fig. 5.**
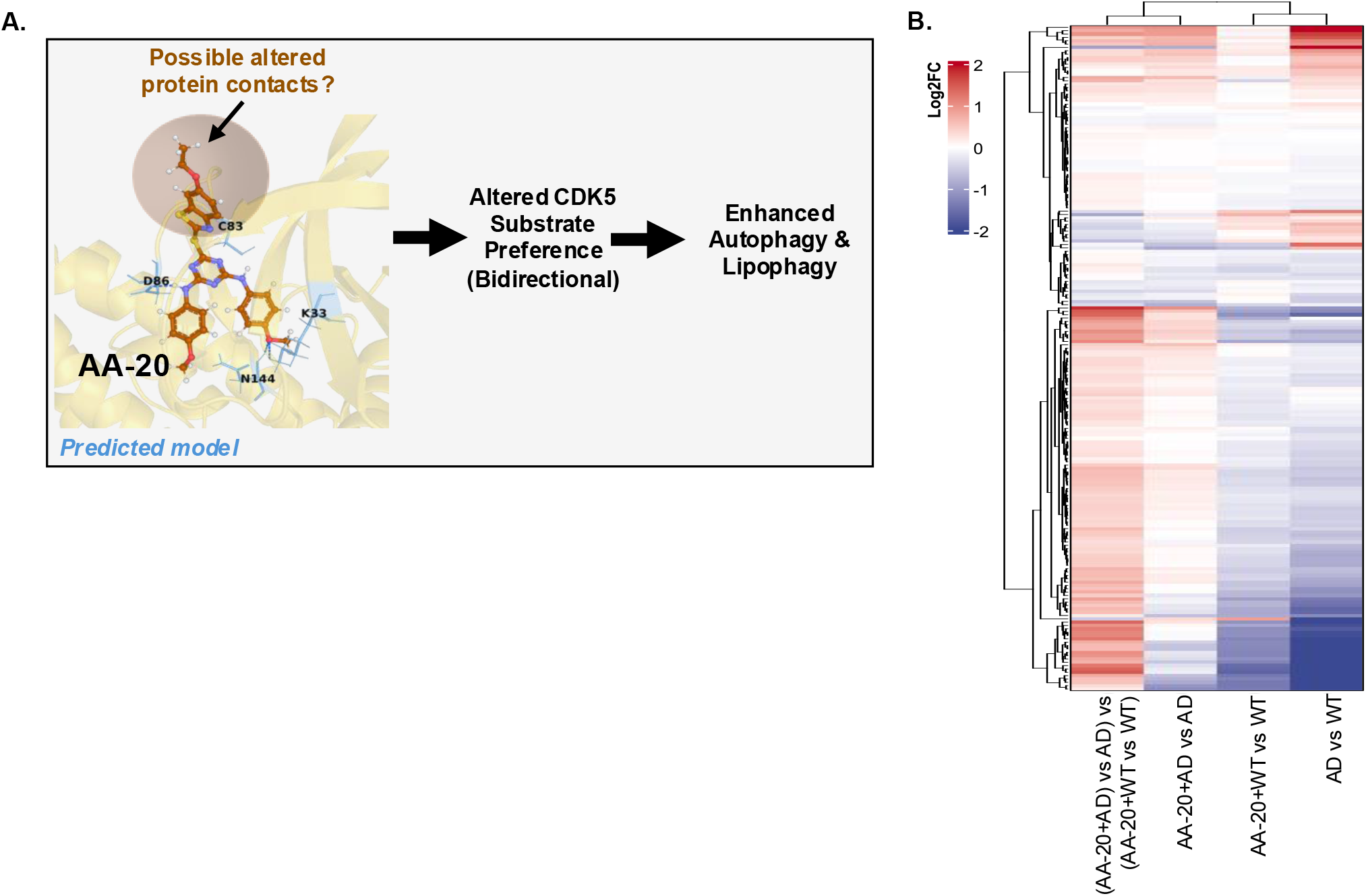
Proposed working model: AA-20 binding to CDK5 may alter substrate preference. (A) Schematic illustrating the proposed mechanism of AA-20 as a CDK5 modulator. AA-20 is predicted to bind the CDK5 ATP-binding pocket without acting as a classical inhibitor. In this working model, AA-20 binding alters CDK5 substrate engagement, producing bidirectional changes in CDK5-dependent phosphorylation and promoting autophagy and lipophagy. This schematic is a predicted model. (B) Heatmap displaying log_2_-fold changes in 199 phosphorylation sites whose parent proteins are targeted by CDK5. Hierarchical clustering was performed using the Euclidean distance and the complete linkage method. Conditions are as follows: “AD”, APPswe/WT; “WT AA-20”, WT AA-20/WT; “AD AA-20”, APPswe AA-20/APPswe; and “AD AA-20 – WT AA-20”, APPswe AA-20/APPswe vs. WT AA-20/WT (see Supplemental Data 12 for complete dataset).

### Chronic AA-20 treatment bidirectionally modulates CDK5 substrate phosphorylation in human iPSC-derived cerebral organoids

The HT22 data indicated that AA-20 binds CDK5 but does not act as a classical inhibitor, raising the possibility that it modulates CDK5-dependent phosphorylation in a context-specific manner (**Fig. 5A**). To test this in a human disease model, we analyzed known CDK5 substrate phosphosites in the same 3-month-old APPswe/WT and isogenic WT organoids described above (**SI Fig. 5A**, **Supplemental Data 2** (for TMT information) using the same statistical contrasts as in the proteomics analysis (**Fig. 2H**).

We identified 23 established CDK5 substrate, encompassing 199 phosphosites. Untreated AD organoids showed a mixed pattern relative to WT: hyperphosphorylation of several neuronal/developmental proteins (e.g., TBR1 for cortical layering; DCX and MAP1B for neuronal migration and cytoskeleton; GPRIN1 for neurite outgrowth; HIVEP2 and SOX5 for transcription; IGF2BP1 for RNA regulation) and hypophosphorylation of multiple cytoskeletal proteins involved in axonal structure and transport (e.g., multiple MAP1B sites; neurofilaments NEFM/NEFH/NEFL/PRPH; doublecortin family DCX/DCLK1/2 for microtubule dynamics) (*65, 80–83*) (AD vs isogenic WT; intense red cluster at top, blue at bottom; **Fig. 5B**; **Supplemental Data 12**). The above described 14 day AA-20 treatment protocol reversed many of these changes in AD organoids (AA-20+AD vs AD), with minimal effect in WT organoids (AA-20+WT vs WT; **Fig. 5B**; **Supplemental Data 12**). The interaction contrast ((AA-20+AD vs AD) – (AA-20+WT vs WT)) indicated that this normalization was stronger in the AD background than in WT, with AA-20 increasing phosphorylation at many hypo-sites and decreasing phosphorylation at many hyper-sites selectively in the AD organoids (e.g., neuronal migration, cytoskeletal integrity, cortical layering (*65, 80–83*) (**Fig. 5B**; **Supplemental Data 12**).

These results indicate that AA-20 bidirectionally and disease-selectively normalize phosphorylation of known CDK5 substrates in AD organoids. Together with the HT22 binding and functional data, this supports a model in which AA-20 acts as a CDK5 modulator rather than a simple activator or inhibitor (**Fig. 5A**).

### AA-20 reduces pathological aSyn-phosphorylation in a neuronal seeding model of PD

Because AA-20 improved AD-related phenotypes in cerebral organoids, we next asked whether AA-20 could also ameliorate PD-relevant proteinopathy. Lewy bodies (LB) contain aggregated aSyn together with membrane, vesicles, and lipid-rich material, and aSyn-lipid interactions have been implicated in fibril formation and seeding (*24, 36, 37*). We therefore tested AA-20 in a well-established primary-neuron seeding neuronal model of aSyn fibrillization and LB-like inclusion formation (*35, 84*) (**Fig. 6A**). In this model, preformed aSyn fibrils (PFFs) are taken up by hippocampal neurons and template recruitment of endogenous aSyn, producing neuritic and somatic aggregates. Recombinant mouse PFFs were characterized by SDS-PAGE, Thioflavin T fluorescence, and negative-stain EM before use (**SI 6A-D**). Pathology was quantified by phosphorylation at Ser129 (pS129-aSyn), a post-translational modification that is low under physiological conditions but highly enriched in pathological aSyn aggregates in PD (*85*).

**Fig. 6.**
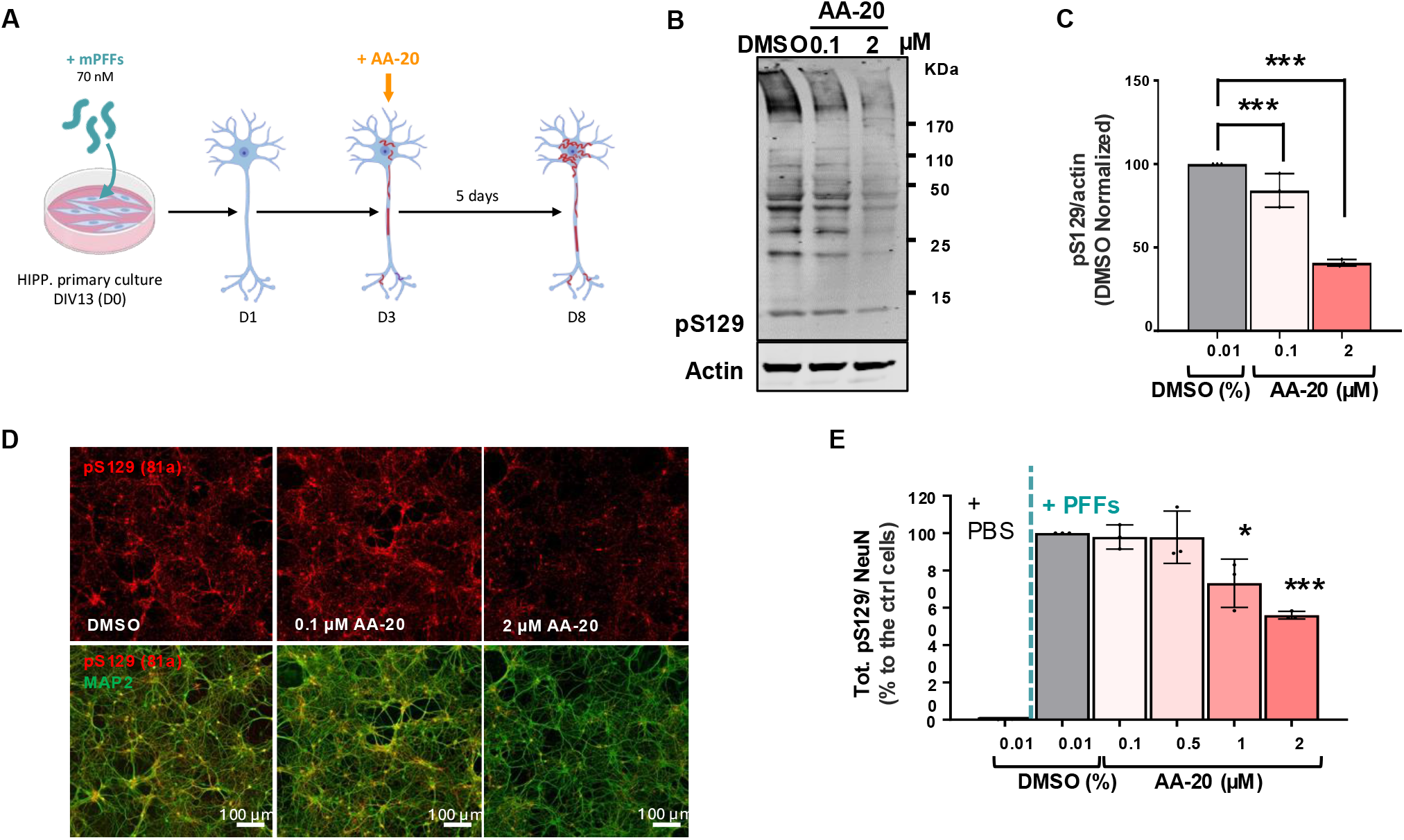
AA-20 reduces pathological aSyn phosphorylation in a neuronal seeding model of PD. (A) Schematic illustration of the seeding model in neurons. Briefly, at day in vitro (DIV) 13, PBS (negative control) or mouse aSyn PFFs were added at 70 nM to WT hippocampal primary cultures. 3 days later, AA-20 or DMSO (0.01%) was added to the PFF-treated cultures. 5 days after compound addition, primary cultures were lysed and analyzed by IB (B-C) or fixed and analyzed by ICC (D-G). (B) Total cell lysates of PFF-treated neurons, treated for 5 days with DMSO or AA-20 at a final concentration of 0.1 or 2 µM, were analyzed by IB. (C) Immunoblotting with anti-pS129 aSyn confirmed a significant reduction in pS129 levels upon AA-20 treatment. pS129 levels were normalized to actin. (D) Representative images of the pS129-aSyn-positive seeded aggregates formed in neurons. Primary cultures were fixed and immunostained with an antibody against pS129 pathology (81a pS129 antibody) (red). Neurons were counterstained with the MAP2- and NeuN-specific neuronal markers (green). Scale bar = 100 µm. (E) The total level of pS129-aSyn-positive seeded aggregates formed in neurons. All the quantifications are based on a minimum of three independent biological replicates, each derived from a separate primary neuronal culture preparation. For HCA analyses, data were normalized to the number of neurons in each acquired field of view and expressed as a percentage reduction relative to PFF-treated neurons incubated with DMSO (control condition). For each biological replicate, three independent wells were analyzed per condition, and nine non-overlapping fields of view were acquired per well. Error bars represent the standard deviation across biological replicates. Statistical significance was determined using one-way ANOVA followed by Tukey’s HSD post hoc test *\*p*<0.05, *\*\*p*<0.01, *\*\*\*p*<0.001, *\*\*\*\*p*<0.0001.

We first added AA-20 (0.1–2 µM) on day 3 (D3) after PFF seeding, when pS129-aSyn aggregates begin to form, and analyzed cultures on day 8 (D8) (**Fig. 6A**). IB showed a ∼60% reduction in pS129-aSyn when AA-20 was employed at a 2 µM concentration, and a ∼20% reduction when used at 0.1 µM (**Fig. 6B-C**). High-content imaging analysis (HCA) with a pS129-aSyn antibody (81a) and the neuronal marker MAP2 (labeling cell bodies and neurites) showed a dose-dependent decrease in total pS129-aSyn after normalization per neuron, with ∼35% reduction at 1 µM and ∼45% reduction at 2 µM of AA-20 (**Fig. 6D-E**). The reduction was greater in neurites (50%) than in cell bodies (∼35%) at 2 µM (**Fig. 6E, SI Fig. 6E**), indicating that AA-20 more strongly attenuates neuritic than somatic aSyn pathology. The pS129-aSyn signal was normalized per neuron. Co-treatment of DIV13 (13 days *in vitro*) neurons with AA-20 and PFFs produced a similar reduction (**SI Fig. 6F-I**), indicating that AA-20 was effective both at the start of seeding and after early aggregates had begun to form.

Seeding efficiency depends on three key factors: 1) PFF uptake, 2) endogenous aSyn levels, and 3) cellular proteostasis. To test whether AA-20 reduced aSyn pS129 by altering PFF internalization or processing, we used aSyn KO primary neuronal cultures, which cannot form seeded aggregates and therefore allow PFF fate to be tracked independently (*35*). After co-treatment with 70 nM PFF and AA-20 (0.1–2 µM), IB showed no change relative to DMSO in SDS-resistant high-molecular-weight PFF levels at 6 h, 24 h (D1), and 48 h (D2), in C-terminal 12 kDa cleavage fragments, or in residual PFF levels at day 7 (**SI Fig. 6J**). Thus, AA-20 did not detectably alter PFF uptake, proteolytic processing, or residual PFF levels at day 7.

To determine whether the AA-20 dependent reduction in pS129-aSyn was attributable to neuronal loss, we quantified DAPI+/NeuN+ neurons. At 2 µM, AA-20 reduced neuron count by 20% relative to PFF-treated DMSO controls (**SI Fig. 6K**). At 1 µM, neuron count was unchanged, yet pS129-aSyn load was reduced by 35% (refer to **Fig. 6D-E**). At 2 µM, pathology decreased by 45-50% (see **Fig. 6D-E**) while neuron count decreased by only 20% (**SI Fig. 6K**). Thus, the reduction in pS129-aSyn cannot be explained by neuronal loss alone. Together, these data show that AA-20 reduces established and newly seeded aSyn pathology in primary neurons without detectably changing PFF uptake or processing.

## DISCUSSION

AA-20 enhances lipid clearance in neuronal systems, including under nutrient-replete conditions and in a human AD organoid model. In HT22 cells under nutrient-rich conditions, AA-20 reduced oleic acid-induced LDs more effectively than Torin 1 and increased association of LDs with autophagic compartments by EM analysis. In human iPSC-derived AD cerebral organoids, 14 days of AA-20 treatment reduced disease-associated elevations in triacylglycerides, diacylglycerols, fatty acids, and cholesteryl esters in an LAL-dependent manner. Proteomics profiling showed that chronic AA-20 treatment preferentially upregulated lipid-clearance enzymes, including LIPA/LAL, in the AD background rather than broadly suppressing biosynthetic pathways. These results indicate that AA-20 can correct selected features of lipid dysregulation in an AD model, particularly when lipid-clearance pathways are already altered.

A second major finding is that AA-20 binds CDK5 and does not behave as a classical CDK5 inhibitor. Direct binding was supported by NanoBRET, CETSA, and SPR assays. Docking and MD simulations indicated that AA-20 can occupy the CDK5 ATP-binding pocket in a pose overlapping that of Roscovitine, yet AA-20 only weakly inhibited CDK5/p35 activity toward histone H1. One possible explanation is that AA-20 modulates CDK5 activity or substrate engagement that is consistent with a molecular glue-like mechanism rather than blocking catalysis outright. Whether AA-20 alters CDK5 protein-protein interactions remains speculative and will require direct testing. The large 7.6 °C T_m_ increase produced by AA-20 treatment suggests that a CDK5-involved protein-protein interaction may be involved in the mechanism of AA-20 mediated autolysosomal degradation. In AD organoids, AA-20 bidirectionally shifted phosphorylation of known CDK5 substrates toward the WT pattern, with little effect in isogenic WT organoids. This disease-selective phosphoproteomic profile is consistent with modulator-like behavior, but it does not by itself demonstrate that all of AA-20’s effects are CDK5-dependent.

We therefore propose, rather than conclude, a working model in which AA-20-bound CDK5 contributes to ULK1-dependent autophagy and lipophagy. CDK5 is enriched in neurons and has been implicated in cytoskeletal regulation, intracellular trafficking (*65, 80–83*), and autophagy initiation (*32, 33*). In HT22 cells, AA-20 increased ULK1 phosphorylation at Ser777 (AMPK site) and Ser622 in a CDK5/2-dependent manner. Inhibition of CDK5/2 partially blocked and ULK1 mostly blocked AA-20-induced autophagic flux, CTSD activation, and LD reduction. These data support CDK5/2-ULK1 signaling as one pathway contributing to AA-20-mediated autophagy and LD clearance in this cell model. Whether CDK5 directly phosphorylates ULK1, and whether CDK5 modulation specifically facilitates LD recruitment, remain to be determined.

AA-20 also reduced pS129-aSyn pathology in a PFF-seeded primary-neuron model of PD, without detectably changing PFF uptake or processing. This experiment did not measure LDs or lipophagy. A more direct interpretation is that enhanced lysosomal function could influence aSyn turnover, as pathological aSyn species are known to be degraded in part through lysosomal pathways (*86, 87*). Consistent with this, reduced activity of the lysosomal enzyme glucocerebrosidase (GCase) is a major genetic risk factor for PD is linked to impaired lysosomal function and aSyn accumulation (*88*). Lipid-rich membranes and LDs have also been associated with aSyn aggregation (*24, 36, 37*), but the present data do not establish that AA-20 reduces aSyn pathology by clearing LDs.

The disease-selective effects observed in AD organoids and the reduction in pS129-aSyn pathology in the PFF-seeded primary-neuron model of PD provide a rationale for further evaluation, but the therapeutic implications are unclear. AA-20 produced minimal apparent perturbation of CDK5 substrate phosphorylation in WT organoids, and improved selected lipidomic, phosphoproteomic, and hyperexcitability phenotypes in the AD background. Whether the reductionin hyperexcitability depends on lipophagy was not tested. These findings are consistent with our previous work showing that AA-20 extends *C. elegans* lifespan through autophagy- and TFEB-related pathways (*15*). Possible links among CDK5, ULK1, and TFEB (*89, 90*) are all interesting but remain inferential.

Taken together, these data identify AA-20 as an mTORC1-independent compound that can promote lipophagic lipid clearance in neuronal cells and correct lipid abnormalities in human AD organoids in a LAL-dependent fashion. Collectively, the CDK5 binding, disease-selective CDK5-substrate remodeling, and reduced seeded aSyn pathology in primary neurons support AA-20 as a candidate neuronal lipophagy-modulating compound. In particular, LAL-dependent lipid clearance by an mTORC1-independent autophagy activator has been little explored in neuronal cells and human brain organoids. Establishing whether lipophagy is the primary mechanism in the organoid and aSyn models, and whether these effects hold in vivo, will be important next steps.

## AUTHOR CONTRIBUTION

Conceptualization: EPT, DR, DG, EP, MP, MH, SF, MAP, HAL, JWK

Methodology: EPT, LA, ALMM, YZ, SRL, SL, AM, DR, JAP

Investigation: EPT, LA, ALMM, YZ, PMB, SRL, LAM, ZG, ZA, KN, CA, WR, CC, YW, JC, MA, OL, CAC, MAP

Funding acquisition: EPT, HAL, JWK

Supervision: EPT, DF, SPG, MP, SAL, SF, MAP, HAL, JWK

Writing – original draft: EPT, LA, ALMM, YZ

Writing – review & editing: EPT, MP, SAL, MH, SF, MAP, HAL, JWK

## COMPETING INTEREST

Prof. Hilal A. Lashuel is the founder and chief scientific officer of ND BioSciences, Epalinges, Switzerland, a company that develops diagnostics and treatments for neurodegenerative diseases (NDs) based on platforms that reproduce the complexity and diversity of proteins implicated in NDs and their pathologies. Scripps Research has filed a patent on AA-20, but a company has not been formed to develop AA-20 into a clinical candidate. Prof. Stuart A. Lipton (S.A.L) is an inventor on worldwide patents for memantine, NitroSynapsin, and related compounds that decrease electrical hyperexcitability in Alzheimer’s and NRF2 transcriptional activators for neurodegenerative and neurodevelopmental disorders. He participates in a royalty-sharing agreement with his former institution, Boston Children’s Hospital/Harvard Medical School, which licensed the FDA-approved drug memantine (Namenda®) to Forest Laboratories, Inc./Actavis/Allergan/AbbVie. S.A.L. is also a scientific founder of Adamas Pharmaceuticals, Inc. (now owned by Supernus Pharmaceuticals, Inc.), which developed or comarketed FDA-approved forms of memantine- or amantadine-containing drugs (NamendaXR®, Namzaric®, and GoCovri®), and of EuMentis Therapeutics, Inc., (which licensed NitroSynapsin). He also serves on the Scientific Advisory Board of Point6 Bio. The rest of the authors declare no competing interests.

## ACKNOWLEDGMENTS

We thank Drs. Matthew Holcomb, Manuel Llanos, and Diogo Santos-Martins for the help and expertise they provided with the computational simulations. We also thank Sara Cano-France for technical support, Yllza Jasiqi and Dr. Theodora Panagaki for their help with the preparation of the primary neuronal cultures used in this work and characterization of the mouse WT PFF used in this study. We thank Dr. Fabien Kuttler (SV PTECH PTCB, EPFL) for his tremendous help in developing the HCA pipeline of analysis. We thank Nicholas McKee for generating the lipophagy schematic. Expert writing and editing support was provided by Dr. Emily P. Bentley. This work was supported in part by CIRM grant EDUC4-12811 and National Institute on Aging (NIA) grant K99 AG080109 and 4R00AG080109-03 to E.T.; Ecole Polytechnique Fédérale de Lausanne (Switzerland) and the Fondation Bru to A.L.M; Endowed Fellowship in the Skaggs Graduate School of Chemical and Biological Sciences to Y.Z.; NIA AG067331 to M.P; NIA AG038664 to M.H.; NIA RF1AG073418 and P01AG054407, and Freedom Together Foundation to J.W.K.. NIA R35 AG071734, R01 AG077046 and U01 AG088679 to S.A.L.; and by Project CP21/00017 funded by the Instituto de Salud Carlos III (ISCIII) and co-funded by the European Union and Project PID2022-136403OA-I00 funded by MCIN/AEI/10.13039/501100011033/FEDER, UE to M.A.P. The content of this article is solely the responsibility of the authors and does not necessarily represent the official views of the National Institutes of Health.

## METHODS

### Statistical Analysis

Electron microscopy and confocal microscopy data were analyzed by one-way analysis of variance (ANOVA). Immunoblot densitometry analyses and flow cytometry analyses were done by one-way ANOVA or two-way ANOVA with Tukey’s multiple comparisons test, respectively. One-way ANOVA with Tukey’s Honest Significant Difference post hoc test was used for all hippocampal primary neuron experiments. The statistical tests used are stated in each figure legend. *p* < 0.05 was considered statistically significant.

### Cell Culture

HT22 mouse hippocampal neuronal cells were a gift from the Michael Petrascheck lab, which he received as a gift from Pam Maher. HT22-GFP-LC3-RFP-LC3ΔG engineered cell line was generated by retroviral transduction of a plasmid containing a GFP-LC3-RFP-LC3ΔG construct. The retrovirus was generated using the pMRX-IP-GFP-LC3-RFP-LC3ΔG plasmid (Addgene #84572) through the Sanford Burnham Prebys Shared Resource Viral Vector Service. HT22 cells were infected with the retrovirus for 48 h, followed by one-week of puromycin selection. The resulting stable reporter cell line was passaged before use. HT22 cell lines were maintained by culture at 37°C and 5% CO_2_ in Dulbecco’s Modified Eagle Medium (DMEM) supplemented with 10% fetal bovine serum (FBS, Omega Scientific) and 1% Penicillin-Streptomycin (Gibco).

### Transmission Electron Microscopy

HT22 cells (250-300K) were plated onto a 10 cm cell culture dish in supplemented DMEM and pre-treated with 60 μM OA-BSA (Sigma, O3008) for 20 h. Cells were treated with 0.2% vehicle control (DMSO), autophagy-activator positive control Torin 1 (150 nM, LC Laboratories T-7887), and AA-20 (20 μM) for 24 h. Cells were then washed with serum-free DMEM and immediately fixed with oxygenated 4% PFA and 2.5% glutaraldehyde in 0.1M sodium cacodylate buffer (pH 7.39) for 1 h on ice. Samples were then washed and fixed in buffered 1% osmium tetroxide + 1.5% potassium ferrocyanide for 1 h on ice and stained overnight at 4 °C with 0.5% uranyl acetate. Samples were dehydrated with a graded ethanol series and subsequently with acetone, fixed in epoxy resin (Ladd Research #21210), and polymerized at 60 °C. Ultrathin sections (60 nm) were obtained using a UC7 ultramicrotome (Leica) and imaged at 80 kV with a Talos L120C transmission electron microscope (ThermoFisher). Images were taken with a CETA 16-M CMOS camera (ThermoFisher); 50 cells per treatment condition were counted to determine the mean number of electron-dense degradative compartments per cell. Additionally, the number of degradative compartments per cell containing lipid droplets was counted.

### Preparation and treatment of aSyn KO and WT hippocampal primary neurons

Primary hippocampal neurons were obtained from postnatal day 0–1 (P0–P1) mouse pups derived from either wild-type (WT) C57BL/6J backgrounds or aSyn knockout (aSyn KO; C57BL/6JOlaHsd) mice as previously described (*35*). WT animals were sourced from Janvier Labs (France), whereas aSyn KO mice were bred in-house at the EPFL animal facility. Neonatal pups were euthanized by decapitation immediately prior to tissue collection, and all experiments were performed ex vivo. Animal procedures were conducted in accordance with institutional and national guidelines and were approved by the Swiss Federal Veterinary Office (VD3694).

Under sterile conditions, hippocampi were rapidly dissected in ice-cold Hank’s Balanced Salt Solution (HBSS) and enzymatically dissociated using papain (20 U/mL) for 30 min at 37 °C. Enzymatic activity was stopped by adding a protease inhibitor solution, followed by gentle mechanical trituration to obtain a single-cell suspension. Cells were collected by centrifugation, resuspended in Neurobasal medium supplemented with B27, L-glutamine, and penicillin/streptomycin, and plated onto poly-L-lysine–coated culture vessels. Neurons were seeded in either black clear-bottom 96-well plates for high-content imaging analyses or in 6-well plates for biochemical assays at a final density of 300,000 cells/mL. Cultures were maintained at 37 °C in a humidified incubator with 5% CO₂ and left undisturbed except for experimental manipulations.

### PFFs preparation, characterization and neuronal treatment

Recombinant mouse aSyn PFFs were prepared by dissolving 4 mg of purified, lyophilized full-length aSyn in 600 µl sterile PBS (pH 7.4), filtering the solution through a 0.22-µm centrifugal filter and incubating it in sealed low-binding tubes for 5 days at 37 °C and 1,000 rpm as previously described (*35*). Fibrils were recovered by centrifugation at 100,000g for 30 min at 4 °C, resuspended in 600 µl PBS and sonicated on ice using a UP200St fine probe at 40% amplitude with 1-s ON/1-s OFF pulses for 10 s, repeated five times with cooling between cycles. Each preparation was characterized by ThT fluorescence, SDS–PAGE and Coomassie staining after separation of the fibrillar, oligomeric and monomeric fractions, and negative-stain EM; accepted batches contained ≤5% soluble monomeric and oligomeric aSyn and predominantly 50–100-nm fibrils after sonication (*91*). Validated PFFs were divided into 15-µl aliquots, snap-frozen in liquid nitrogen and stored at –80 °C.

One aliquot of PFFs was thawed immediately before treatment of primary neuronal cultures. After DIV13, WT or aSyn KO neuronal cultures were exposed to extracellular mouse WT aSyn PFF at a final concentration of 70 nM to induce seeded aggregation (*35*). Control cultures received an equivalent volume of PBS. Small-molecule treatments were performed by adding the AA-20 compound, prepared as a 100 µM or 1 mM stock solution in DMSO, directly to the cultures at final concentrations ranging from 0.1 to 20 µM. Vehicle controls received DMSO alone at a final concentration of 0.01%. Neurons were maintained under these conditions until the experimental endpoint.

### Immunofluorescence and confocal microscopy

HT22 cells (15K) were seeded onto 8-well confocal plates (ibidi, 80826) in supplemented DMEM. Cells were incubated at 37 °C for 3 h to allow full adherence, then pre-treated with 60 μM OA-BSA and 0.1% DMSO, 150 nM Torin 1, and 20 μM AA-20 the following day for 24 h. Cells were fixed with 4% paraformaldehyde (PFA) in PBS at room temperature (RT) for 20 min, then stained with LipidTOX (1:250, Invitrogen, H34475) or BODIPY (1:1000, Invitrogen, D3922) for neutral lipids and Hoechst nuclear dye (1:1000, ThermoFisher Scientific, 62249). For DAPRed/LipidTOX colocalization experiments, cells were stained with DAPRed (1:200, Dojindo, D677) prior to fixation. Imaging was performed using a Zeiss LSM 880 confocal scanning microscope. Total LD area and DAPRed-LipidTOX colocalization by Manders coefficient were quantified on Imaris.

For primary neuronal pS129/NeuN/MAP2 IF and quantitative high-content analysis (HCA), we used previously described methods (*35*). Briefly, at the experimental endpoint, primary hippocampal neurons treated with mouse aSyn PFF in the presence or absence of AA-20 were rinsed twice with PBS and fixed for 20 min at RT using 4% PFA. Fixed cultures were permeabilized and blocked according to standard protocols before incubation with primary antibodies. Aggregated aSyn was visualized using a phosphorylation-dependent antibody recognizing serine 129 (pS129; clone 81a), which preferentially labels pathological aSyn species. Neuronal architecture (soma and neuritic extensions) were identified by immunolabeling for microtubule-associated protein 2 (MAP2). Neuronal nuclei were detected using NeuN immunostaining, and all nuclei present in the mixed neuronal–glial cultures were labeled with DAPI. Antibody sources and dilution factors are specified in the corresponding Table 1 in Supplementary Information.

**Table 1:** List of the primary and secondary antibodies used.

| Antibodies | Catalog# | Company | Host | Concentration | WB dilution | ICC dilution |
|---|---|---|---|---|---|---|
| aSyn total | 610787 | BD | Mouse | 0.25 mg/mL | 1:1000 | 1:1000 |
| pS129-asyn | 825701 | BioLegend | Mouse | 1 mg/mL | 1:1000 | 1:2000 |
| pS129-asyn | Ab168381 | Abcam | Rabbit | 4.18 mg/mL | 1:10 000 | n/a |
| MAP2 | Ab92434 | Abcam | Chicken | Not provided | n/a | 1:2000 |
| NeuN | Ab177487 | Abcam | Rabbit | 1 mg/mL | n/a | 1:5000 |
| Actin | Ab6276 | Abcam | Mouse | 2.2 mg/mL | 1:5000 | n/a |
| Mouse Alexa Fluor 680 | 926-68070 | Li-Cor | Goat | 1 mg/mL | 1:20 000 | n/a |
| Rabbit Alexa Fluor 800 | 926-32211 | Li-Cor | Goat | 1 mg/mL | 1:20 000 | n/a |
| Mouse Alexa Fluor 647 | A31571 | Invitrogen | Donkey | 2 mg/mL | n/a | 1:10 000 |
| Rabbit Alexa 568 | A10042 | Invitrogen | Donkey | 2 mg/mL | n/a | 1:10 000 |
| Chicken Alexa Fluor 488 | 703-545-155 | Jackon Immunoresearch | Donkey | 1 mg/mL | n/a | 1:10 000 |
| LAL | Sc-58374 | Santa Cruz | Mouse | 0.2 mg/mL | 1:1000 | n/a |
| ULK1 | A7481 | Millipore Sigma | Rabbit | 1 mg/mL | 1:1000 | n/a |
| pS757-ULK1 | CST6888S | Cell Signaling Technology | Rabbit | 1 mg/mL | 1:1000 | n/a |
| pS777-ULK1 | ABC213 | Sigma-Aldrich | Rabbit | 1 mg/mL | 1:1000 | n/a |
| pS623-ULK1 | AB248371 | Abcam | Rabbit | 1 mg/mL | 1:1000 | n/a |
| P70-S6K | CST2708S | Cell Signaling Technology | Rabbit | 1 mg/mL | 1:1000 | n/a |
| PS6K-T389 | CST9206S | Cell Signaling Technology | Mouse | 1 mg/mL | 1:1000 | n/a |
| CDK5 | CST12134S | Cell Signaling Technology | Mouse | 1 mg/mL | 1:1000 | n/a |
| Tubulin | CST8058S | Cell Signaling Technology | Mouse | 1 mg/mL | 1:1000 | n/a |

Images were taken on an automated wide-field high-content imaging platform (IN Cell Analyzer 2200) using the Nikon 10×/0.45, Plan Apo, CFI/60 of the IN Cell Analyzer 2200 (GE Healthcare), a high-throughput imaging system equipped with a high-resolution 16-bit sCMOS camera (2048×2048 pixels)) (*35, 92*). All images were collected using identical acquisition parameters to ensure comparability across experimental conditions.

Quantitative image analysis was carried out using a predefined segmentation workflow (*35*). This multiparametric approach enabled compartment-resolved quantification of pS129-positive aSyn pathology within neuronal cell bodies and MAP2-defined neuritic processes (see Table 1 for the antibodies references). In parallel, neuronal survival (NeuN⁺/DAPI⁺ nuclei), neuritic network integrity (MAP2-positive area), and non-neuronal cell populations (DAPI⁺/NeuN⁻ nuclei) were quantified. Collectively, these measurements allowed the effects of AA-20 on aSyn aggregation to be distinguished from changes in neuronal viability or overall culture composition.

### Immunoblotting

For HT22 cells, cells were seeded onto 6-well plates and treated with 60 μM OA-BSA and small molecules for 20-24 h. Cells were then washed with PBS, dissociated with 0.25% trypsin-EDTA (Gibco), pelleted and lysed on ice with lysis buffer containing 1% NP-40, 150 mM NaCl, 50 mM Tris-HCL, pH 7.5, phosphatase inhibitor and protease inhibitor for approximately 30 min, vortexing every 5 min. Lysates were clarified by centrifugation for 5 min (maximum speed, 4 °C), then boiled in Laemmli buffer at 95 °C for 5 min; the protein-containing supernatant was normalized by BCA using the Pierce BCA Protein Assay Kit (Thermo Fisher). Proteins were separated by SDS-PAGE (4-15% gradient gels, Bio-Rad, #4561086) and transferred onto a PVDF membrane (Bio-Rad Trans-Blot Turbo). Membranes were blocked with 5% bovine serum albumin (BSA) in Tris-buffered saline with 0.05% Tween-20 (TBST) and labeled with primary antibodies overnight at 4 °C (see Table 1 for the complete list of antibodies). Membranes were washed 3x with TBST and probed with horseradish peroxidase (HRP)-conjugated anti-mouse or anti-rabbit secondary antibodies (Cell Signaling Technology) for 1 h at RT. Blots were washed at least 3x with TBST before imaging by chemiluminescence (Bio-Rad ChemiDoc Imaging System).

For total cell lysates preparation and WB analysis of aSyn KO and WT hippocampal primary neurons, PBS- or PFF-treated aSyn KO and WT neurons in which AA-20 or DMSO compounds were added for up to 8 days were lysed in 2% SDS, 50 mM Tris, 150 mM NaCl, pH 7.4 supplemented with protease inhibitor cocktail (Sigma-Aldrich), phenylmethane sulfonyl fluoride (PMSF; Applichem), and phosphatase inhibitor cocktails 2 and 3 (Sigma-Aldrich). The total cell lysate was boiled for 10 mins. The protein concentration in each sample was quantified using the Bicinchoninic acid protein assay (Thermofisher) before the addition of Laemmli buffer 4x (10% SDS, 50% glycerol, 0.05% bromophenol blue, 1M Tris-HCl pH 6.8 and 20% β-mercaptoethanol). Proteins were next separated on 1-mm-thick 16% Tricine gels (*93*) for 2 h at 120 V. The proteins were transferred to 0.2 µm nitrocellulose membranes (Amersham) using a semi-dry transfer system (Life Technologies) under constant current (0.5 A) and a voltage of 25 V. The membranes were blocked for 30 min at RT in Intercept blocking buffer (Li-COR Biosciences) and probed with the primary antibodies overnight at 4°C (see table in the “Antibodies” section of materials and methods). The membranes were then washed three times with PBS containing 0.1% (V/V) Tween 20 (Fluka) before being incubated with secondary goat anti-mouse or anti-rabbit antibodies conjugated to Alexa Fluor 680 or 800 (1:20,000; Li-COR Biosciences). Then, three washes with PBS containing 0.01% (V/V) Tween-20 were performed. The membranes were finally scanned on the Li-COR scanner (Li-COR Biosciences). The levels of total aSyn, pS129 aSyn, LC3 and p62 were quantified using Image Studio software (*RRID*:SCR_015795; Li-COR Biosciences), and normalized to the relative protein levels of actin.

### Flow Cytometry

HT22 cells were seeded onto 6-well or 96-well plates in supplemented DMEM and incubated at 37°C to allow for adherence, then pre-treated with 60 μM OA-BSA overnight (16-20 h). Media were replaced with 0.1-0.2% DMSO- or AA-20-containing DMEM for 24 h. To measure the requirements of LD clearance pharmacologically, AA-20 was co-treated with Lalistat 2 (10 μM, 6099, Tocris Bioscience), CDK5 inhibitors 20-223 (5 μM, MedChem Express, HY-123772), or ULK-101 (5 μM, MedChem Express, HY-114490). Neutral lipids were then labeled with LipidTOX Green neutral lipid stain (1:250, 1:500 or 1:1000) or BODIPY (1:1000, Invitrogen, D3922) for 30-45 min. Cells were lifted with 0.25% trypsin-EDTA and resuspended in supplemented DMEM before centrifugation to a cell pellet, then washed with PBS before a second centrifugation. Cells were resuspended in stain buffer (BD Bioscience, 554656) and measured on the Bio-Rad Yeti ZE5 Analyzer at 488 nm. Data are represented as LipidTOX or BODIPY mean intensity.

To measure autophagic flux by flow cytometry, we utilized an engineered HT22-GFP-LC3-RFP-LC3ΔG reporter cell line. GFP-LC3 degradation by autophagy is measured as a marker for autophagic flux by changes in GFP fluorescence, while RFP-LC3ΔG, unable to undergo lysosomal degradation due to deletion of the C-terminal glycine, serves as an internal control (developed by Mizushima et al., https://doi.org/10.1016/j.molcel.2016.09.037). Cells were seeded on a 96-well plate and treated with vehicle (DMSO) or AA-20 (1 μM, 10 μM, or 20 μM) for 24 h. To measure AA-20’s dependence on CDK5 for autophagy activation, AA-20 was treated at four doses (1.25 μM, 2.5 μM, 5 μM, 10 μM) alone or with co-treatment of 5 μM 20-223 for 24 h. Cells were prepared as described above, and fluorescence was measured by flow cytometry at 488 nm (GFP) and 561 nm (RFP). Data are represented as GFP/RFP ratio.

### Lysosomal Acid Lipase Activity Assay

200K HT22 cells were seeded into 6-well plates in supplemented DMEM and incubated to allow for full adherence. Cells were then treated with vehicle (DMSO), and AA-20 (10 μM or 20 μM) for ∼22 h. The following was performed according to the LysoLive Lysosomal Acid Lipase Assay Kit (Abcam, ab253380). Growth medium was aspirated and replaced with LysoLive LipaGreen stain in serum-free DMEM (1:1000) for 3 h. After staining, cells were dissociated using 0.25% trypsin-EDTA, resuspended in serum-free DMEM and centrifuged to form a pellet. DMEM was aspirated and replaced with 1X Flow Holding and Sorting Buffer for Adherent Cells (from kit). Cells were measured for mean intensity by flow cytometry on the Bio-Rad Yeti ZE5 Analyzer using the 488 nm laser. Data were analyzed by one-way ANOVA.

### Cellular Thermal Shift Assay (CETSA)

The published protocol was followed with minor modifications (https://doi.org/10.1038/nprot.2014.138). HT22 cells were seeded 24 h before treatment and harvested using 0.25% trypsin-EDTA. Cell pellets were washed twice with ice-cold PBS and resuspended in lysis buffer (1× PBS, 1 mM MgCl₂, 0.5% NP-40, phosphatase inhibitors, and protease inhibitors, pH 7.4). After 10 min of incubation on ice, lysates were centrifuged at 21,000 × g for 30 min at 4 °C. The supernatant was collected, and protein concentration was quantified using a BCA assay, then normalized to 2 mg/mL with lysis buffer. Samples were incubated with DMSO or 100 μM AA-20 for 30 min at RT, aliquoted into 200 μL PCR tubes, and subjected to heat treatment in a thermal cycler with temperature gradients (37–53°C: 37, 40.6, 43.3, 47, 49.8, 53°C; 56–67°C: 56, 58.6, 62.9, 67°C) for 3 min. A control sample was maintained on ice. After cooling to RT, samples were ultracentrifuged at 100,000 × g for 20 min at 4 °C, and the supernatant was collected. Approximately 25 μg of protein per condition was resolved by SDS-PAGE and transferred to PVDF membranes. Membranes were blocked with 5% BSA in TBST and probed overnight at 4°C with a primary antibody against CDK5 (Cell Signaling Technology, 12134S). After washing with TBST, membranes were incubated with HRP-conjugated secondary antibodies (Cell Signaling Technology) for 1 h at RT. Blots were washed three times with TBST and visualized using chemiluminescence.

### Cathepsin D Activity Assay

Cathepsin D activity was assessed using the BioVision Cathepsin D Activity Fluorometric Assay Kit (#K143-100) in HT22 cells, following methods adapted from Tan et al. (*15*). Briefly, HT22 cells were seeded at 2 × 10^5^ cells/well in 6-well plates and incubated overnight in DMEM with 10% FBS. Cells were then treated with DMEM containing 10% FBS and either 0.1% DMSO (vehicle control), 10 μM or 20 μM AA-20 (as specified in figure legends), 250 nM Torin 1, 150 nM BafA1, or co-treated with 5 μM 20-223 or 5 μM ULK-101. After 24 h, cells were washed once with ice-cold PBS, lysed with buffer (100 mM NaAc, 100 mM NaCl, 1 mM EDTA, 1% NP-40, 10 mM DTT, pH 5.5), and scraped from wells. Lysates were centrifuged at 13,000 rpm for 5 min at 4 °C, and supernatants were collected in fresh tubes. Total protein was quantified using the Pierce 660 nm Protein Assay. For the assay, 0.25 μg of total protein per sample was loaded into microplate wells, with samples run in technical triplicate. Fluorescence was measured every min for 30 min at 25 °C using a BMG LabTech PHERAStar FS microplate reader (Ex: 328 nm, Em: 460 nm). Cathepsin D activity was determined by linear regression analysis in GraphPad Prism to calculate the rate of fluorescent substrate cleavage.

### NanoBRET Binding Assay

NanoBRET™ assays (Promega) were conducted by Promega Corporation. Briefly, HEK293 cells were seeded at 7,000 cells/well in 384-well plates, transfected with 2 μg/ml DNA (1:9 kinase: carrier) using FuGENE HD in Opti-MEM with 1% FBS, and incubated overnight at 37 °C. Test and control compounds (1000X) were serially diluted 1:3 in DMSO (10-point series), transferred with 1X tracer and DMSO to an Echo source plate, and dispensed into assay plates. For lytic assays, 10 μl of 3X NanoBRET™ Nano-Glo® Solution (1:166 in Opti-MEM, no serum/phenol red, with 1X digitonin) was added post-treatment, incubated for 20 min at RT, and emissions were measured at 450 nm (donor) and 610/630 nm (acceptor) using a NanoBRET™ luminometer.

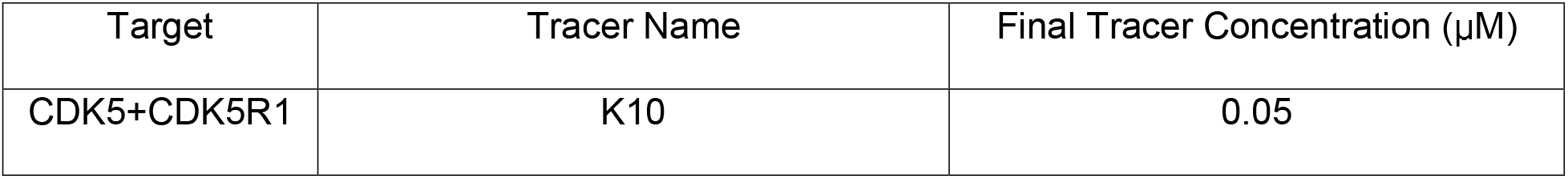

| Target | Tracer Name | Final Tracer Concentration (µM) |
|---|---|---|
| CDK5+CDK5R1 | K10 | 0.05 |

For live-cell assays, 10 μl of 3X NanoBRET™ Solution (with 1:500 extracellular inhibitor) was added after 2-h treatment and 15-min equilibration, followed by 3-min incubation and emission measurement.

| Target | Tracer Name | Final Tracer Concentration (µM) |
|---|---|---|
| CDK5+CDK5R1 | K10 | 1 |

Raw BRET ratios (acceptor/donor) were converted to milliBRET (mBU) by multiplying by 1000, and IC50 values were determined using JMP with a 4-parameter Hill curve.

### AutoDock Vina Docking and Molecular Dynamics (MD) Simulation

The crystal structure of the CDK5^D144N^/p25 complex (PDB ID: 1UNL, https://www.ncbi.nlm.nih.gov/Structure/pdb/1UNL) was used as the receptor for molecular docking and MD simulations. The CDK5/p35 complex structure is not available due to crystallization challenges, so p25 was used instead. The D144N mutation was introduced to improve expression yields. Water molecules and heteroatoms were removed, and missing atoms were added. Hydrogen atoms were added, and side chain orientations of Asn, Gln, and His were optimized using the Computational Crystallography Toolbox (cctbx, v2025.6)(*94*). The 3D structure of AA-20 was generated from its SMILES representation using molScrub (https://github.com/forlilab/molscrub). Both the ligand and receptor were converted to PDBQT format using Meeko (v0.6.1) (https://github.com/forlilab/Meeko), which was also used to define a docking box encompassing the entire protein to allow unbiased sampling of potential binding sites. Molecular docking was performed using AutoDock Vina (v1.2.7)(*70*). 50 independent docking runs were conducted with an exhaustiveness value of 10, each generating the top 9 binding poses, yielding a total of 450 conformations. The resulting poses were clustered based on spatial proximity based on RMSD-based hierarchical clustering.

MD simulations were performed using OpenMM (v8.1.1)(*95*). The protein was modeled with the AMBER ff14SB force field, and the ligand was parameterized using the OpenFF 2.0.0 small molecule force field. The system was solvated in a cubic TIP3P water box with 1.0 nm padding, and Na^+^/Cl^-^ ions were added to a final concentration of 0.15 M. A 1.0 nm cutoff was used for nonbonded interactions. Equilibration included gradual heating from 100 K to 300 K, followed by multi-stage positional restraint release on the protein backbone, side chains, and ligand. All trajectory frames were aligned to a reference structure using protein backbone heavy atoms with MDTraj (*96*) to remove overall rotation and translation prior to analysis.

Two types of MD simulations were performed. Standard MD simulations were run starting from the top-ranked docking pose of AA-20 in the NPT ensemble at 300 K for 200 ns using a 2 fs timestep. Three independent replicates were performed. Trajectory frames and state data were saved every 25,000 steps for analysis. RMSD and RMSF calculations were performed using MDAnalysis (v2.9.0)(*97, 98*) on the protein backbone and ligand heavy atoms. Steered MD simulations were carried out for two ligand-protein complexes: AA-20 bound in the ATP-binding pocket from a representative frame at the end of standard MD, and Roscovitine in its crystallographic binding pose (PDB ID: 1UNL, https://www.ncbi.nlm.nih.gov/Structure/pdb/1UNL). For each complex, three replicates were run using the same equilibrated structure but different initial velocity seeds. Each simulation was 1.5 ns at 300 K. A harmonic restraint with a force constant of 2000 kJ/mol/nm^2^ was applied between the center of mass of the ligand and that of the protein backbone, and the restraint target was displaced by 1.2 nm at constant velocity. Trajectory frames and simulation states were saved every 500 steps.

Other analytical software packages used include ProDy (*99*), SciPy (*100*), NumPy (*101*), pandas (*102*), and Matplotlib (*103*). All structural visualizations were created using PyMOL (*104*).

### Human iPSC cultures

All experiments involving human cells were conducted with Institutional Review Board approval at The Scripps Research Institute and as described previously (*52, 105, 106*). Two sets of human induced pluripotent stem cells (hiPSCs) were utilized: the previously generated and well-characterized isogenic heterozygous APPSwe/WT mutant line, and its wild-type parental counterpart (WT 7889SA) (*50*). Both hiPSC lines were cultured concurrently on vitronectin-coated plates (Thermo Fisher), using mTeSR Plus medium (StemCell Technologies), which was refreshed daily. Cells were visually inspected during routine maintenance; any with abnormal morphology were manually removed when necessary. Passaging was performed every 5–6 days using the ReleaSR reagent (Thermo Fisher), enabling gentle dissociation of cell colonies without enzymes. Mycoplasma contamination was regularly monitored, and cells were not propagated beyond 13 passages before replacement.

### Cerebrocortical Organoid Cultures

Cerebral organoids were generated using a recently developed protocol in our laboratory, similar to Paşca et al. (*52, 107*). Briefly, hiPSCs at 75–90% confluence were dissociated with Accutase and seeded as single cells (4 × 10^6^ per well) into Aggrewell plates in mTeSR Plus medium with CEPT cocktail: 50 nM Chroman I, 5 μM Emricasan, 1× Polyamine, and 0.7 μM Trans-ISRIB. Embryoid bodies (EBs) formed over 24 h at 37 °C and were then transferred to ultra-low-attachment 6-well plates on a 3.5°-tilted orbital shaker (60 rpm) in a 1:1 mix of Essential 6 and hESC medium, supplemented with 2.5 μM Dorsomorphin, 10 μM SB-431542, and 2.5 μM XAV-939 for 6 days. The medium was then switched to Neural Media-A containing 20 ng/ml EGF2 and FGF2. From days 25–43, these supplements were replaced by 20 ng/ml BDNF and NT-3, after which organoids were maintained in Neural Media-A. Around day 35, organoids were transferred to ultra-low attachment 10 cm dishes and cultured on a 3.5°-tilted shaker. Organoids were monitored, split as needed, and tested for mycoplasma every 1–2 months.

### Microelectrode Array (MEA) Recordings

Cerebral organoids (COs, 6–8 weeks old) were plated individually onto 48-well MEA plates (Axion Biosystems) pre-coated overnight at 37 °C with 0.05% poly(ethyleneimine) (Sigma-Aldrich) followed by laminin I (R&D Systems) in PBS. After removing the laminin solution, COs were positioned at the well center, allowed to attach for 15 min at 37 °C, and wells were then filled with BrainPhys medium supplemented with NeuroCult SM1 and N2 Max. Medium was partially changed every two days, and COs were acclimated for at least one week prior to recording. Electrophysiological data were collected for 5-min epochs 23–24 h after the last medium change, at consistent daily timepoints, using the Axion Maestro Pro system, as we have previously described in detail (*105, 106*). Data were analyzed using Axion’s recommended protocols in the Navigator app, with parameter extraction via Axion’s Neurometric Tool app.

### Cerebral Organoid Treatments

Individual cerebral organoids were transferred to separate wells of ultra-low attachment 24-well plates, each containing a standardized media volume (e.g., 1.5 mL). DMSO was added to ensure a consistent vehicle concentration across all experimental conditions. Treatments were administered with each media change (every two days), while plates were agitated on orbital shakers at 100 rpm. For MEA electrophysiology experiments, baseline recordings were collected for at least one week prior to compound exposure. At the start of treatment, compounds were freshly diluted in BrainPhys medium, with concentrations adjusted for partial media exchange. Throughout the experiment, organoid viability was monitored in real time by electrode resistance measurements at each recording session. Mean activity for each cerebral organoid was calculated from multiple technical replicates.

### Lipidomic Analysis by Mass Spectrometry (MS)

Human iPSC-derived organoids were treated with either DMSO (vehicle control), AA-20 (5 μM), or AA-20 in combination with the lysosomal acid lipase inhibitor Lalistat 2 (10 μM; Tocris Bioscience, #6099) for approximately 13 days. Following treatment, organoids were harvested, washed with ice-cold PBS to remove residual media, and stored at –80 °C until lipid extraction and mass spectrometry analysis.

For extraction, organoids were washed with DPBS and transferred to pre-weighed Eppendorf tubes, then centrifuged at 500 × g for 4 min. The supernatant was removed, and the resulting pellet was lyophilized overnight. Samples were normalized based on the dry weight of the lightest sample by adding 400 μL/mg of ddH₂O containing 1 mM erythorbate. A total of 500 μg of sample was homogenized on ice using a 22-gauge needle (15 passes). To avoid plastic contamination, all subsequent steps were performed using glass pipettes and tubes.

Total lipids were extracted by adding 400 μL/mg of cold chloroform (CHCl₃) containing a 1:50 dilution of SPLASH® Lipidomix® internal standards (Avanti Polar Lipids). Samples were vortexed and centrifuged in glass test tubes at 200 × g for 5 min. The bottom organic phase was collected using a glass Pasteur pipette, and the upper aqueous phase was re-extracted twice with solvent mixtures in the following order: H₂O:MeOH:CHCl₃ (1:2:2, v/v), followed by MeOH:CHCl₃ (2:1, v/v). A final extraction was performed using MTBE:H₂O (4:1, v/v), and the resulting organic layers were pooled and dried under vacuum to yield a lipid film, which was stored at –80 °C until analysis.

Dried lipid films and external standards (SPLASH® Lipidomix®, Avanti) were resuspended in 100 μL of methanol. Positive and negative ion mode LC-MS was performed using an Agilent 6230 ESI-TOF-MS system, calibrated with reference masses at 922 m/z and 1033.988 m/z. Separation was achieved on an XBridge BEH C8 XP Column (Waters; 2.5 µm, 4.6 mm × 150 mm) at a flow rate of 500 μL/min. The LC gradient consisted of 40–100% solvent B over 40 min, with the first 10 min held at 40% isocratic B. Solvent A was composed of Milli-Q water and methanol (1:1, v/v), and Solvent B consisted of MTBE:2-propanol (1:4, v/v); both contained 2 mM ammonium acetate or 2 mM sodium acetate.

Raw data files (.d format) were converted to the open-format mzXML using MSConvert (ProteoWizard toolkit; Chambers et al., 2012). Mass feature detection was performed using the wavelet algorithm in MZmine 2, with ADAP chromatogram builder and deconvolution (Myers et al., 2017), all within the MZmine 3 software package (Pluskal et al., 2010). Lipid identifications were performed using the LIPID MAPS database with a mass accuracy threshold of 15 ppm.

### Quantitative tandem mass tag (TMT) proteomics and phosphoproteomics

HT22 cells were seeded in 10 cm dishes (n = 16) and cultured in supplemented DMEM for 24 h prior to treatment with the indicated compounds for an additional 24 h. After treatment, cells were washed three times with ice-cold PBS, harvested by scraping into PBS, and cell pellets were stored at –80°C until further processing. Organoids were cultured and harvested as previously indicated, in DMSO (vehicle control) or AA-20 (5 μM), for approximately 13 days. Whole proteome and phosphoproteome analyses were performed according to our previously published method (*108*) and based on the streamlined tandem mass tag (SL-TMT) workflow developed by the Gygi laboratory (*109*). For lysis, cells were resuspended in 8 M urea lysis buffer (8 M urea, 75 mM NaCl and 50 mM HEPES, pH 7.5) supplemented with EDTA-free protease and phosphatase inhibitors (Roche). Lysates were then clarified by centrifugation at 17,000 x g for 15 min at 4 °C, and protein concentrations were determined using a bicinchoninic acid (BCA) assay (Thermo Fisher Scientific).

For each sample, 200 µg of total protein was reduced with 5 mM tris(2-carboxyethyl)phosphine (TCEP) for 30 min, alkylated with 14 mM iodoacetamide for 30 min in the dark, and then treated with 10 mM dithiothreitol (DTT) for 15 min, all at RT. Next, proteins were precipitated using the chloroform-methanol method, and the resulting protein pellets were resuspended in 200 mM EPPS (pH 8.5). Samples were then digested overnight at 37 °C with a mixture of Trypsin/LysC (Thermo Fisher Scientific) at a 1:100 (protease:protein) ratio using an orbital shaker set to 1,500 rpm. After digestion, samples were clarified by centrifugation at 17,000 x g for 10 min, and peptide concentration was measured using the Pierce Quantitative Peptide Assay (Thermo Fisher Scientific).

Next, tandem mass tagging (TMT) labeling was performed using the TMTpro-18plex reagents (Thermo Fisher Scientific, Lot. No.: YJ375600 and YJ372607). Briefly, 100 μg of peptides was adjusted to a concentration of 1 μg/μl with 200 mM EPPS (pH 8.5), followed by the addition of acetonitrile (ACN) to a final concentration of 30%. Next, each sample was labeled with 200 μg of the corresponding TMT reagent and incubated for 1 hr at RT. Labeling reactions were quenched with 0.3% hydroxylamine (Sigma-Aldrich) for 15 min at RT. To assess digestion and labeling efficiency and determine the appropriate mixing ratios for equal peptide loading, a ratio check was performed. For this, 1.5 µl of each sample was mixed and acidified with 3% formic acid (FA) to bring the final ACN concentration to 3%. The samples were then desalted using C18 Stop-and-Go Extraction Tips (STAGE-Tip) (*110*), dried using a SpeedVac concentrator, and subsequently analyzed on an Orbitrap Exploris 480 mass spectrometer (Thermo Fisher Scientific). Based on these analyses, mixing volumes were adjusted, samples were pooled into a final solution containing 3% ACN and 3% FA, desalted using 100 mg tC18 SepPak cartridges (Waters)(*109*), and dried in a SpeedVac.

Next, desalted peptides were subjected to a phosphopeptide enrichment (‘mini-phos’) using the High-Select Fe-NTA Phosphopeptide Enrichment Kit (Thermo Fisher Scientific, Cat. No. A32992) following the manufacturer’s instructions. The flow-through and first two washes were combined, concentrated to near dryness using a SpeedVac, resuspended in 10 mM ammonium bicarbonate (NH_4_HCO_3_), and fractionated by basic pH reversed-phase high-performance liquid chromatography (BPRP-HPLC) using an Agilent Zorbax 300 Extended-C18 column (3.5 μm, 2.1 mm ID × 25 cm). Mobile phases consisted of Buffer A (5% ACN, 10 mM NH₄HCO₃, pH 8) and Buffer B (95% ACN, 10 mM NH₄HCO₃, pH 8). Fractions were first collected into 96-well plates and then combined into 24 fractions, from which twelve nonadjacent fractions were selected, desalted using the STAGE-Tip method and dried down in a SpeedVac before mass spectrometry analysis. These samples correspond to the total proteome analysis. Phosphopeptides eluted from the ‘mini-phos’ enrichment were also desalted using STAGE-Tips and dried down in the SpeedVac before mass spectrometry analysis. Additional sample details are provided in **Supplementary Data 2**.

### Mass spectrometry analysis

All samples were analyzed on an Orbitrap Exploris 480 mass spectrometer operating in high-resolution MS2 (hrMS2) mode (Thermo Fisher Scientific) equipped with a FAIMSpro module and coupled to an EASY-nLC 1200 ultra-high-performance liquid chromatography (UHPLC) system (Thermo Fisher Scientific) (*111, 112*). Peptides were separated on a 100 μm inner diameter microcapillary column packed with ∼35 cm of Accucore C18 resin (2.6 μm, 150 Å, Thermo Fisher Scientific) at a flow rate of 300 nl/min. The mobile phases consisted of Buffer A (5% ACN, 0.1% FA) and Buffer B (95% ACN, 0.1% FA). Detailed parameters for each TMT analysis, including gradient length, FAIMS compensation voltages (CV), MS1 resolution, scan range, MS1 maximum injection time, automatic gain control (AGC), isolation window, higher-energy collision dissociation (HCD), MS2 resolution, MS2 maximum injection time, and AGC, are provided in **Supplementary Data 2.**

### Proteomic data processing and analysis

RAW mass spectra data were converted to .mzML format using MSConvert (ProteoWizard software) (*113*) and then searched using MSFragger (v4.1) (*114*) against a mouse (UP000000589, last modified 2024-06-11; 17,207 reviewed, canonical proteins) or human (UP000005640, retrieved 2024-06-11, containing 20,420 reviewed, canonical entries) target-decoy protein databases that included common contaminants, for the cell line and organoids, respectively. Precursor ion mass tolerance was set at 20 ppm and product ion tolerance at 0.02 Da. TMTpro labels on lysine residues and peptide N-termini (+304.20715 Da) and carbamidomethylation of cysteine residues (+57.021 Da) were set as static modifications, while oxidation of methionine residues (+15.995 Da), and phosphorylation (+79.9663 Da) of serine/threonine/tyrosine residues in phosphoproteomic analyses, were set as a variable modifications. The search was restricted to tryptic and LysC-generated peptides, allowing up to two missed cleavage sites. MSFragger output files were processed using Philosopher (v5.1.1): firstly, PSMs were adjusted to a 1% false discovery rate (FDR) using Percolator (*115*), followed by peptides being assembled into proteins by ProteinProphet (*116*). Protein groups were filtered to 1% FDR using the target-decoy strategy. For all PSMs corresponding to a TMT-labeled peptide, intensities were extracted from the corresponding MS2 scans. The PSM table generated as a Philosopher output file was used by TMT-Integrator (*117*) and was filtered for quantitation quality: minimum peptide probability was set to 0.9, precursor ion purity ≥ 50%, minimum MS1 intensity 0.05%, and summed MS2 intensity ≥ 5%. PSMs mapping to common contaminants were excluded. Lastly, PSMs were aggregated to peptides and proteins. For phosphoproteomic analysis, site probability was set as > 0.75 as an extra filtering step.

All proteomic analyses were conducted in R (v4.4.1) and RStudio (v2024.12.0). Key packages included dplyr (v1.1.4) and tidyr (v1.3.1) for data manipulation, and edgeR (v4.2.2) and limma (v3.60.6) for normalization and data modeling, respectively. Additional Bioconductor packages included clusterProfiler (v4.12.6), org.Hs.eg.db (v3.20.0) and org.Mm.eg.db (v3.19.1) for enrichment analysis, KinSwingR (v1.26.0) for kinase prediction, and ggplot2 (v3.5.1), ggvenn (v0.1.10) and complexHeatmap (v2.20.0) for data visualization.

Data were subject to a two-step normalization to account for both sample loading and compositional biases. Normalized intensities were log2-transformed before being incorporated into the limma framework (*118*) for differential expression analysis. A separate linear model was fitted for each protein using lmFit, comparisons of interest were incorporated via makeContrasts and contrasts.fit, and subsequently fed to eBayes, which incorporated a mean-variance trend. Lastly, topTable was used to calculate the moderated t-statistic and associated p-value for each protein, and those proteins with adjusted p-values < 0.05 (Benjamini-Hochberg [BH] correction to control for the false discovery rate [FDR]) were considered significant.

Gene ontology (GO) enrichment analyses were performed using enrichGO (clusterProfiler), with the set of all proteins quantified in the specific experiment serving as the background, and org.Mm.eg.db and org.Hs.eg.db as the mouse and human, respectively, GO (biological process, molecular function and cellular compartment) database (2024-01). Terms with adjusted p-values < 0.05 (BH correction) were considered significant. Additionally, the rrvgo package was used to reduce GO term redundancy by means of semantic similarity. More specifically, the relevance (“Rel”) method was used to calculate the similarity matrix, which was further reduced using a similarity threshold of 0.9. Parent terms were selected based on adjusted p-values (scores).

Kinase prediction from phosphoproteomic data was conducted using KinSwingR (*119*), which leverages the phosphorylation motif (+/-7 amino acids that surround the phosphorylated S/T/Y residues), the fold change and direction of the change, and the reported p-value of each phosphopeptide. Kinase-substrate information was downloaded from PhosphoSitePlus (v6.8.0)(*120*).

The code for reproducing the analysis and figures related to the proteomic data is freely available at GitHub (github.com/prado-lab/collab-kellylab-aa20-2026). All mass spectrometry proteomic data have been deposited to the ProteomeXchange Consortium via the PRIDE(*121*) partner repository under the dataset identifier PXD085066. The dataset is currently private, the following details allow reviewers to access it in the meantime: Project accession: PXD085066; Token: ZcpHBNTyTdwu.

## Supplemental Information

### SUPPLEMENTAL FIGURE LEGENDS

**Supplemental Fig. 1.**
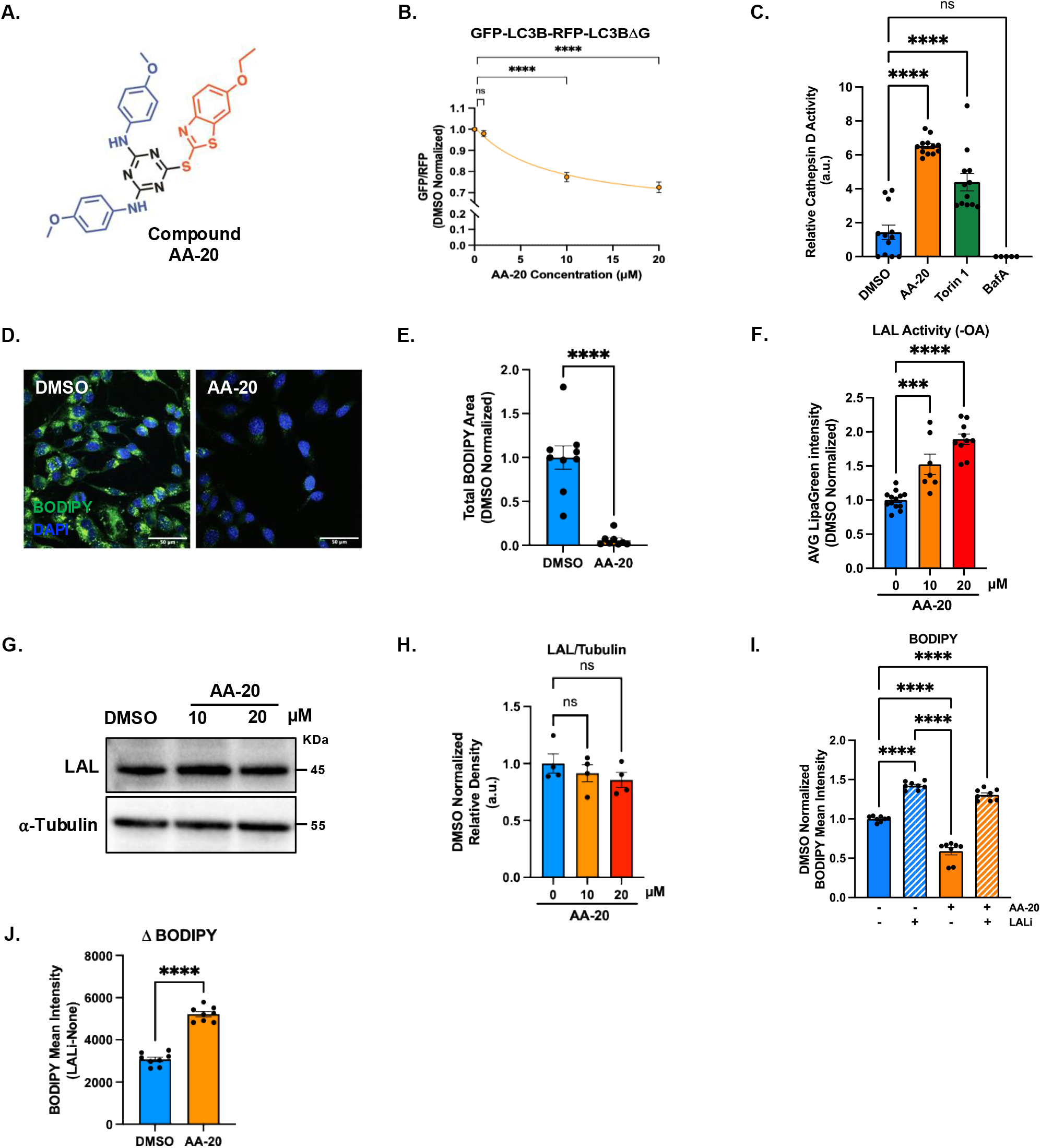
AA-20 acts as an autophagy activator and enhances lipophagy in HT22 cells. (A) Structural representation of the AA-20 compound. Black: 1,3,5-trisubstituted triazine core, blue: 4-methoxyaniline substituents, red: benzothiazole substituent. (B) 3-point dose-response curve of LC3 level by GFP/RFP after 24 h treatment with 1 μM, 10 μM and 20 μM AA-20 in engineered HT22 GFP-LC3-RFP-LC3ΔG reporter cell line, normalized to DMSO. *n=*3 independent experiments. (C) Quantification of CTSD enzymatic activity in HT22 cells after 24 h treatment with either DMSO (0.1%), 10 μM AA-20, 250 nM Torin 1, or 150 nM BafA1. Data represent *N*=12 from six biological replicates, each with two technical replicates. All error bars represent SEM. All statistical significance was determined by ordinary one-way ANOVA (\*\*\*\**p* < 0.0001). (D) Confocal micrographs of HT22 cells stained with BODIPY (green) and Hoechst (blue). Scale bar=50 μm. (E) Bar graph showing data points of total BODIPY area per cell, normalized to their respective DMSO condition per image. HT22 cells were pre-treated with 60 μM of OA-BSA, then treated with DMSO or 10 μM AA-20 for 20 h. Error bars indicate SEM. At least 3 biological replicates, each with 2-3 images taken. \*\*\*\**p* < 0.0001, by unpaired t-test. (F) Quantification of average LipaGreen intensity after 24 h DMSO (0.1%), 10 μM, or 20 μM AA-20 treatment in HT22 WT cells, without OA-BSA pre-treatment, normalized to DMSO. *n=*3 independent experiments, each with 3-4 biological replicates. (G-H) Representative immunoblot (IB) of LAL after 24 h DMSO (0.1%), 10 μM, or 20 μM AA-20 treatment in HT22 cells (H) and its respective densitometry analysis (H). LAL levels were normalized to α-tubulin. *n*=4 biological replicates. All error bars represent SEM. All statistical significance is denoted as not significant (ns); \**p* < 0.05, \*\*\**p* < 0.001, \*\*\*\**p* < 0.0001, by one-way ANOVA. (I) Bar graph showing the mean intensity of BODIPY by flow cytometry in HT22 cells. Cells were preincubated with 60 μM OA-BSA, then treated with either DMSO (0.2%), 10 μM AA-20 alone, or co-treated with 5 μM Lalistat 2, respectively. All error bars represent SEM. All statistical significance is denoted as ns, not significant; \*\**p* < 0.01, \*\*\**p* < 0.001, \*\*\*\**p* < 0.0001, by ordinary one-way ANOVA. (J) Net LAL-dependent change in neutral lipid. Determined by subtracting BODIPY fluorescence intensity from (I) in the presence of Lalistat 2 from that without Lalistat 2 for both DMSO and AA-20 conditions (DMSO+LALi – DMSO (blue bar), AA-20+LALi – AA-20 (orange bar)). Error bars represent SEM. \*\*\*\**p* < 0.0001, by unpaired t-test.

**Supplemental Fig. 2.**
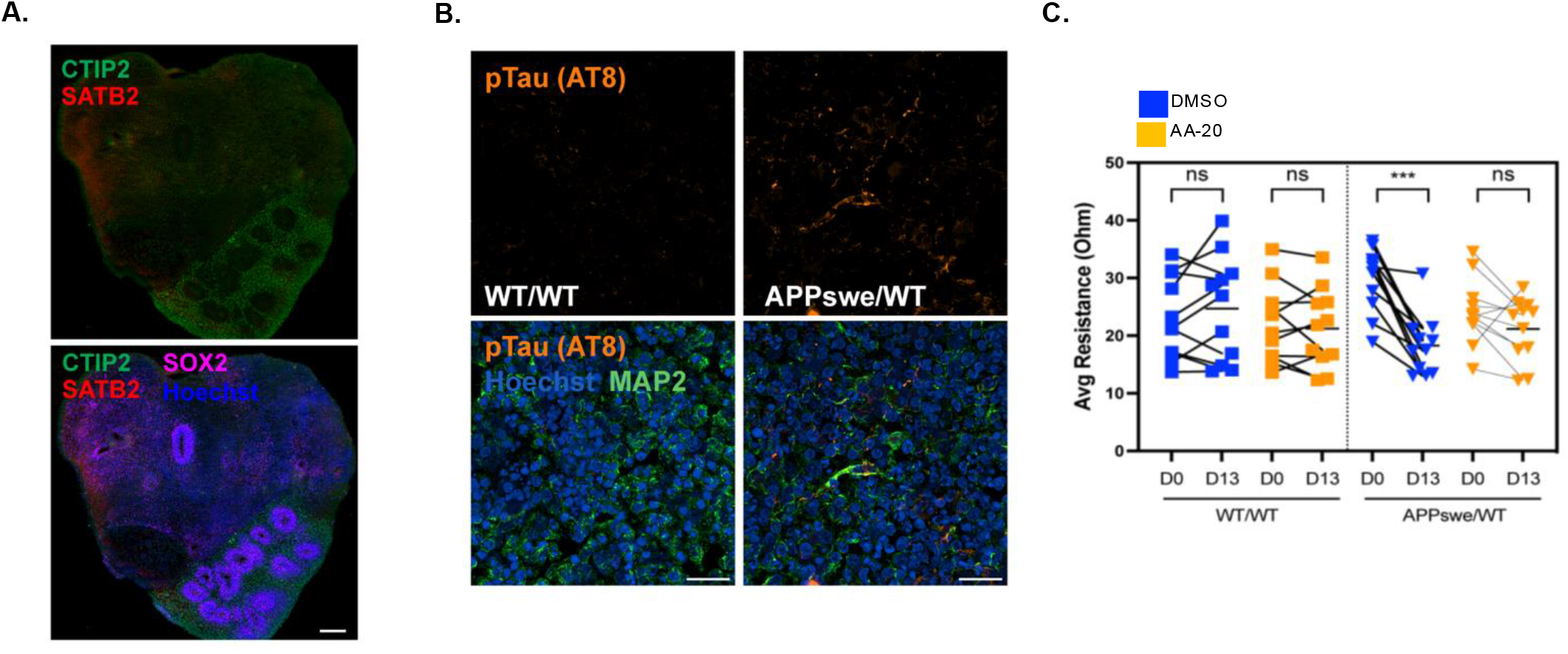
AA-20 normalizes AD-associated pathology in human iPSC-derived organoids. (A) Representative immunofluorescence (IF) image of neuroepithelial progenitors (SOX2^+^) and deep layer V (CTIP2^+^) and upper layer II-III (SATB2^+^) cortical neurons in 3-month timepoint organoids. Scale bar = 200 µm. (B) Representative IF images of phosphorylated Tau fibrils (AT8^+^) and mature cortical neurons (MAP2^+^) in 3-months timepoint organoids. Scale bar = 20 µm. (C) Quantification comparing the organoid impedance (viability index) before (D0) and after the 13-day treatment (D13), measured as average resistance (Ohm). n = at least 10 organoids per condition.

**Supplemental Fig. 3.**
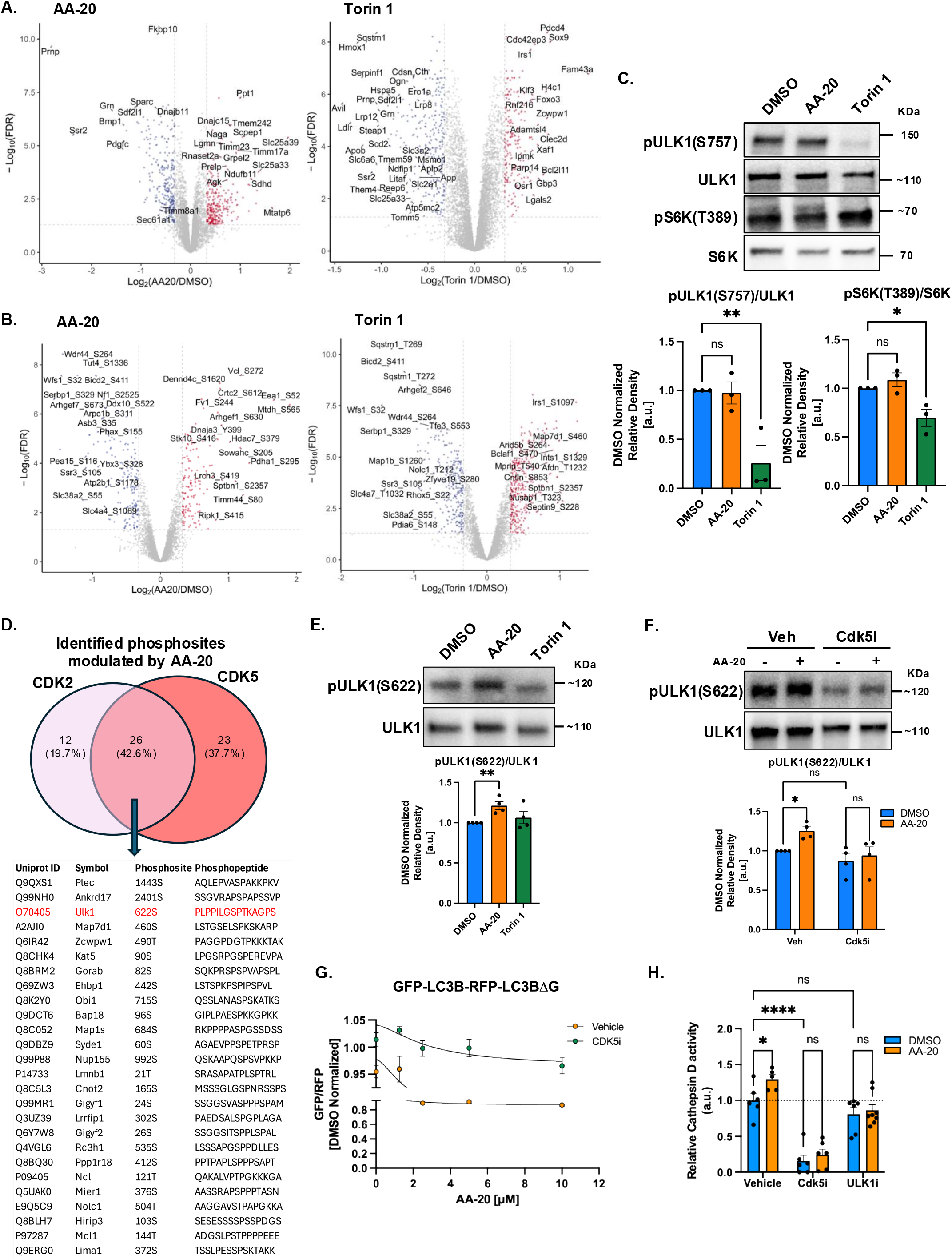
AA-20 distinctly modifies HT22 cell proteomic profiles compared to an mTORC1 inhibitor and promotes ULK1 phosphorylation via CDK5-dependent mechanisms. (A) Volcano plots depicting whole-proteome proteomic analysis of HT22 cells treated with 20 µM AA-20, 150 nM Torin 1, or DMSO, identifying significantly altered proteins relative to DMSO control. AA-20 significantly altered 1,898 proteins, while Torin 1 affected 3,117 proteins. (B) Volcano plots from phosphoproteomic analysis of the same HT22 cell batch as (A), showing significantly altered phosphosites following AA-20, Torin 1, or DMSO treatment. AA-20 significantly modulated 796 phosphosites, and Torin 1 altered 1,525 phosphosites. The x-axis represents log2 fold change between treatments, and the y-axis shows -Log10(adjusted p-value). Grey dotted lines denote thresholds of adjusted p-value < 0.05 (y-axis) and >1.25-fold change (x-axis). Data are based on four biological replicates. Significantly upregulated proteins/phosphosites are marked in red, downregulated ones in blue, with annotations provided (see Supplemental Data 2-3 for complete datasets). (C) Representative IBs and densitometry analyses of pULK1-S757 and pT389-S6K (respectively) after 24 h treatment of DMSO (0.1%), AA-20 (10 µM) and Torin 1 (150 nM). IBs are normalized to total ULK1 or S6K protein levels and the DMSO control. *n=*3 for each phosphorylation site. Statistical significance: ns, not significant; \**p* < 0.05, \*\**p* < 0.01, by ordinary one-way ANOVA. Error bars represent SEM. (D) Venn diagram depicting the number and percentage of phosphosites uniquely and commonly modulated by CDK5 and CDK2, identified via KinSwing analysis of phosphoproteomic data from (B) following AA-20 treatment (see Supplemental Data 6 for CDK5- and CDK2-mediated phosphopeptides). A list of phosphosites and phosphopeptides shared between CDK5 and CDK2 substrates, as determined by KinSwing analysis, is included (see Supplemental Data 7 for complete datasets). (E-F) Representative IBs and densitometry analyses of pULK1-S622, normalized to total ULK1 protein level and DMSO control. (E) HT22 cells were treated with 24 h of DMSO (0.1%), AA-20 (20 µM), or Torin 1 (150 nM), respectively. *n*=4. \*\**p*<0.01, by ordinary one-way ANOVA. (F) Cells were treated with either DMSO or 20 μM AA-20, and co-treated with or without 5 µM 20-223 for 24 h. *n*=4 independent repeats. ns, not significant; \**p* < 0.05, by two-way ANOVA with Tukey’s multiple comparisons. (G) Analysis of GFP/RFP ratio following 4-point AA-20 treatment (1.25, 2.5, 5, 10 µM) with and without CDK5 pharmacological inhibition in HT22 GFP-LC3-RFP-LC3ΔG cells. *n=*3 biological replicates. (H) Quantification of CTSD enzymatic activity in HT22 cells following 24-h treatment with 0.1% DMSO, 20 μM AA-20, or AA-20 co-treated with or without 5 μM 20-223 or 5 μM ULK-101. Data represent three independent experiments, each with at least two biological replicates. Error bars indicate the standard error of the mean (SEM). Statistical significance was determined by two-way ANOVA with Tukey’s multiple comparisons test: ns, not significant; *p < 0.05; ****p < 0.0001.

**Supplemental Fig. 4.**
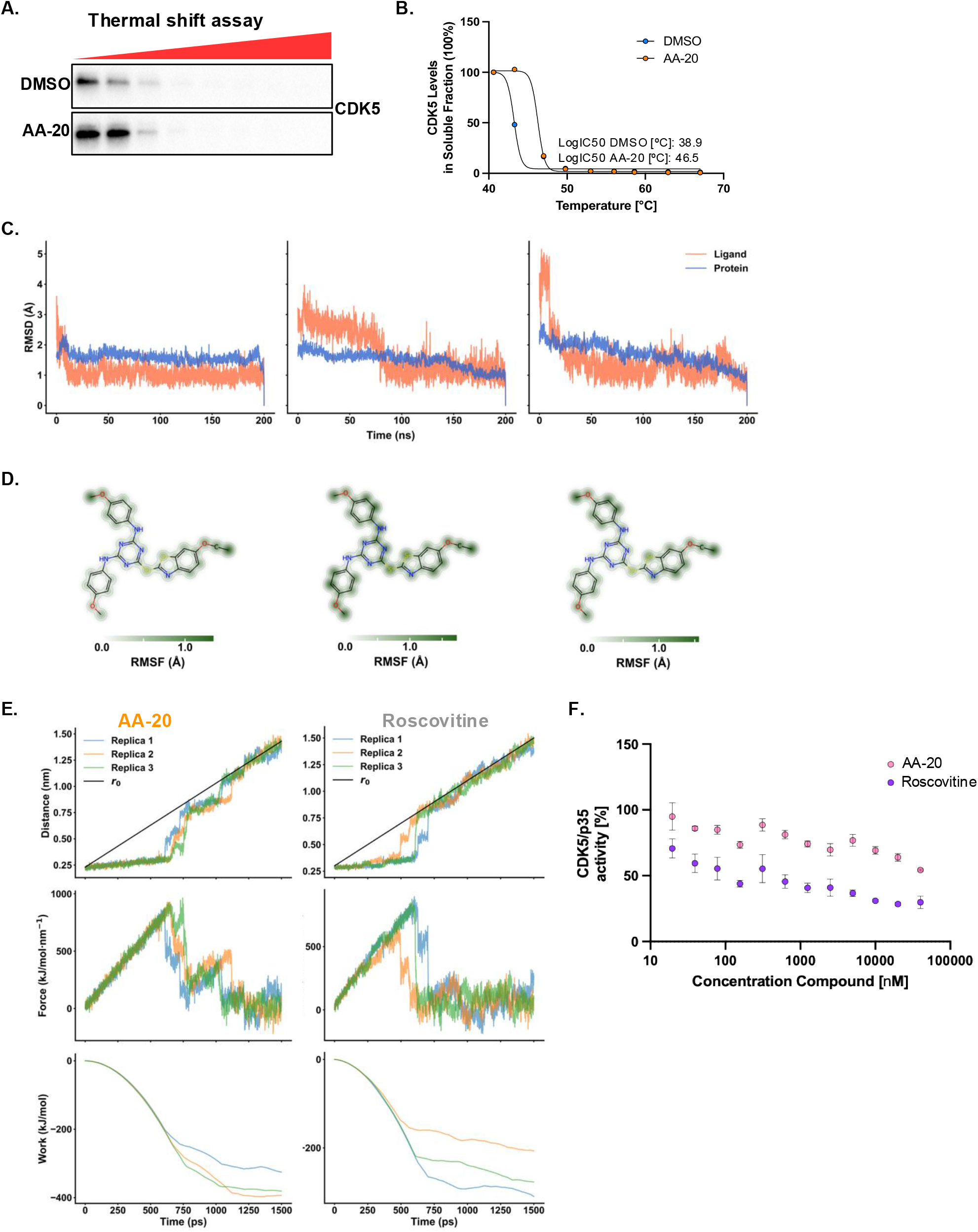
AA-20 acts as a CDK5 modulator. (A) Representative immunoblots of CDK5 following cellular thermal shift assay (CETSA). HT22 WT cells were treated with 0.1% DMSO or 100 μM AA-20 for 30 min, then subjected to gradient heat treatment (37, 40.6, 43.3, 47, 49.8, 53, 56, 58.6, 62.9, 67°C) for 3 min. (B) Densitometry analysis of CDK5 normalized to the 40.6°C treatment for each condition in (A). Data points represent a single replicate per condition. The data were plotted using GraphPad Prism software, and a four-parameter logistic (4PL) regression model was used to generate a sigmoidal curve. *n=*3 independent experiments. (C) RMSD of AA-20 (orange) and CDK5/p25 backbone (blue) over time for each of the three MD replicas. Time (ns) is on the x-axis, RMSD (Å) on the y-axis. (D) Per-atom RMSF of AA-20 mapped onto its structure for each of the three MD replicas. Color intensity reflects atomic fluctuations. (E) Steered MD results for AA-20 (left) and Roscovitine (right), showing ligand-protein center-of-mass distance (top), applied force (middle), and cumulative work (bottom) over time for each of the three simulation replicas. r0 indicates the time-dependent target position of the harmonic restraint. (F) Plot depicting CDK5 kinase activity in response to AA-20 or Roscovitine (CDK5 inhibitor) at concentrations ranging from 0 to 40 μM. In vitro kinase activity was assessed using the ADP-Glo assay (Promega) with recombinant human CDK5/p35 and histone H1 as substrates. Data were analyzed and plotted using GraphPad Prism software, fitting a four-parameter logistic (4PL) regression model to generate sigmoidal dose-response curves. Data represent three independent experiments (n=3), with error bars indicating SEM.

**Supplemental Fig. 5.**
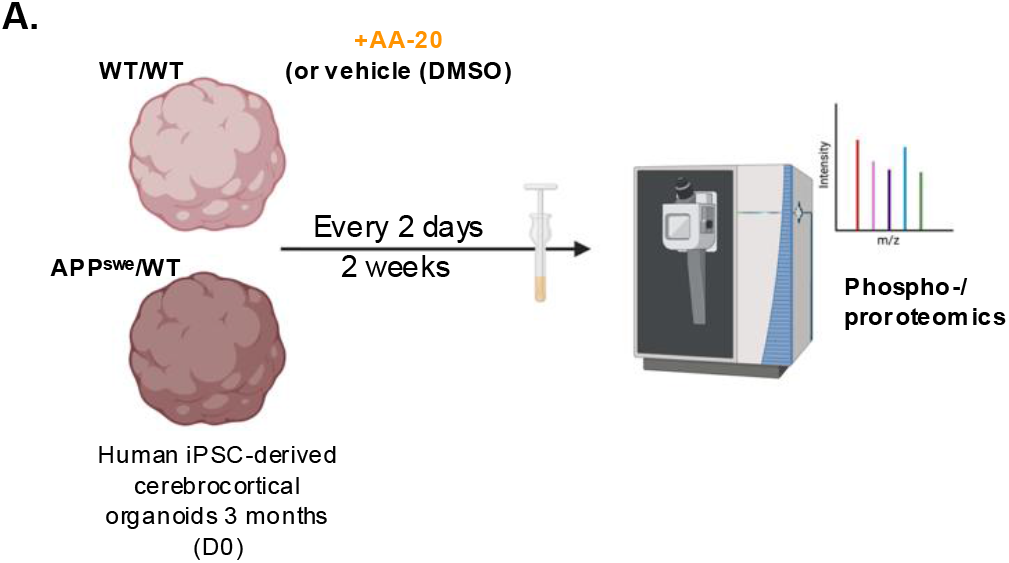
Chronic AA-20 treatment bidirectionally modulates phosphorylation of CDK5 substrates in human iPSC-derived cerebral organoids. (A) Schematic illustrating experimental setup and treatment timeline for human cerebral organoids comparing AD model (APPswe/WT) vs. wild-type control (WT/WT). Created using BioRender.

**Supplemental Fig. 6.**
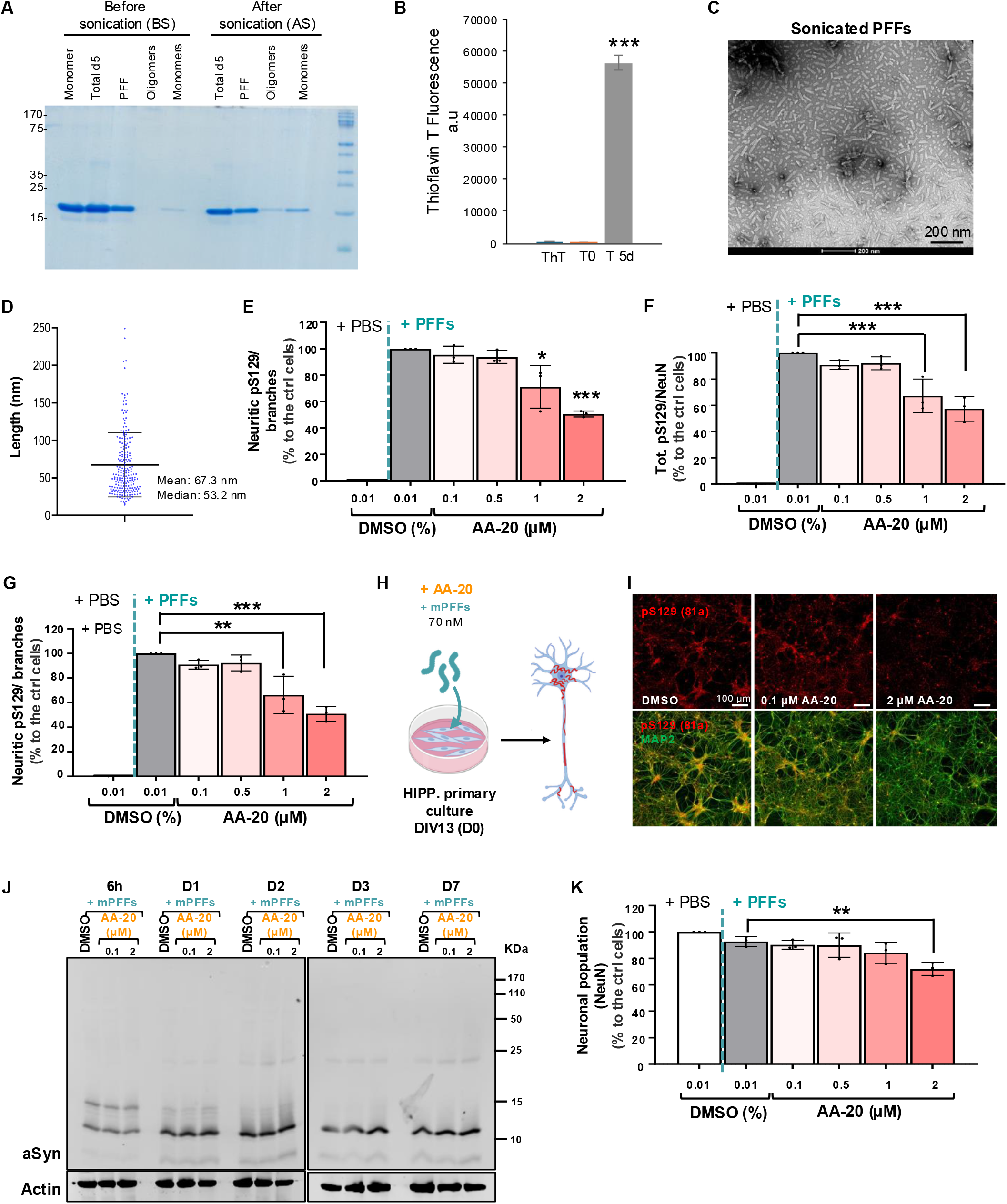
AA-20 did not affect the uptake of PFF in neurons. (A-D) Preparation and characterization of recombinant mouse aSyn PFFs. (A) Coomassie-stained SDS–PAGE of monomeric aSyn at T0 and of the total preparation after 5 days of fibrillization (Total D5), together with the corresponding PFF, oligomeric and monomeric fractions obtained before (BS) and after sonication (AS). (B) ThT fluorescence of ThT alone, monomeric aSyn at T0 and the fibrillized preparation at D5. Statistical significance was assessed using one-way ANOVA followed by Tukey’s post hoc multiple-comparisons test; ***P < 0.001 versus T0. (C) Representative negative-stain EM image of the sonicated PFF preparation. Scale bar, 200 nm. (D) Length distribution of sonicated PFFs measured from EM images. Each point represents one fibril; the mean and median lengths were 67 and 53 nm, respectively. (E) The level of pS129-aSyn-positive seeded aggregates formed specifically in the neuritic branches. (F-G) The total level of pS129-aSyn-positive seeded aggregates formed in neurons (F) and the level of pS129-aSyn-positive seeded aggregates formed specifically in the neuritic branches (G) were measured in the PBS- or PFF-treated neurons incubated with AA-20 (0.1, 0.5, 1 or 2 µM) or with DMSO (negative control). (H) Schematic illustration of the seeding model in neurons. Briefly, at DIV 13, PBS (negative control) or mouse aSyn PFFs at 70 nM were added to WT hippocampal primary cultures with AA-20 or with DMSO (0.01%). After 8 days of co-incubation, primary cultures were fixed and analyzed by ICC combined with HCA. (I) 8 days post-treatment, primary cultures were fixed and immunostained with an antibody against pS129 pathology (81a antibody) (red). Neurons were counterstained with the MAP2-specific neuronal marker (green). Scale bar = 100 µm. (J) Total levels of aSyn and actin were detected by IB. At DIV 13, mouse aSyn PFF were added at 70 nM to aSyn KO hippocampal primary culture with AA-20 at 0.1 µM or 2 µM or with DMSO (0.01%). After 6 h, 1 day (D1), 2 days (D2), 3 days (D3) or 7 days (D7) of treatment, total cell lysates were extracted for IB. (K) The neuronal cell bodies were measured in the PBS- or PFF-treated neurons incubated with AA-20 (0.1, 0.5, 1 or 2 µM) or with DMSO (negative control). All the quantifications are based on a minimum of three independent biological replicates, each derived from a separate primary neuronal culture preparation. For HCA analyses, data were normalized to the number of neurons in each acquired field of view and expressed as a percentage reduction relative to PFF-treated neurons incubated with DMSO (control condition). For each biological replicate, three independent wells were analyzed per condition, and nine non-overlapping fields of view were acquired per well. Error bars represent the standard deviation across biological replicates. Statistical significance was determined using one-way ANOVA followed by Tukey’s HSD post hoc test *\*p*<0.05, *\*\*p*<0.01, *\*\*\*p*<0.001, *\*\*\*\*p*<0.0001.

## Notes

https://github.com/forlilab/molscrub

https://github.com/forlilab/Meeko

https://github.com/prado-lab/collab-kellylab-aa20-2026

## REFERENCES

1. Y. Aman et al., Autophagy in healthy aging and disease. Nat Aging 1, 634–650 (2021).

2. R. Singh et al., Autophagy regulates lipid metabolism. Nature 458, 1131–1135 (2009).

3. J. Chung et al., The Troyer syndrome protein spartin mediates selective autophagy of lipid droplets. Nat Cell Biol 25, 1101–1110 (2023).

4. D. Das et al., VPS4A is the selective receptor for lipophagy in mice and humans. Mol Cell 84, 4436–4453 e4438 (2024).

5. M. Pu et al., ORP8 acts as a lipophagy receptor to mediate lipid droplet turnover. Protein Cell 14, 653–667 (2023).

6. Z. Yuan et al., ATG14 targets lipid droplets and acts as an autophagic receptor for syntaxin18-regulated lipid droplet turnover. Nat Commun 15, 631 (2024).

7. S. Zhang et al., The regulation, function, and role of lipophagy, a form of selective autophagy, in metabolic disorders. Cell Death Dis 13, 132 (2022).

8. E. Turco, D. Fracchiolla, S. Martens, Recruitment and Activation of the ULK1/Atg1 Kinase Complex in Selective Autophagy. J Mol Biol 432, 123–134 (2020).

9. M. Zachari, I. G. Ganley, The mammalian ULK1 complex and autophagy initiation. Essays Biochem 61, 585–596 (2017).

10. J. Kim, M. Kundu, B. Viollet, K. L. Guan, AMPK and mTOR regulate autophagy through direct phosphorylation of Ulk1. Nat Cell Biol 13, 132–141 (2011).

11. R. A. Saxton, D. M. Sabatini, mTOR Signaling in Growth, Metabolism, and Disease. Cell 168, 960–976 (2017).

12. D. W. Lamming, L. Ye, D. M. Sabatini, J. A. Baur, Rapalogs and mTOR inhibitors as anti-aging therapeutics. J Clin Invest 123, 980–989 (2013).

13. S. Sarkar, Regulation of autophagy by mTOR-dependent and mTOR-independent pathways: autophagy dysfunction in neurodegenerative diseases and therapeutic application of autophagy enhancers. Biochem Soc Trans 41, 1103–1130 (2013).

14. L. Yoon, et al., An mTOR-independent Macroautophagy Activator Ameliorates Tauopathy and Prionopathy Neurodegeneration Phenotypes. bioRxiv, 2022.2009.2029.509997 (2023).

15. E. P. Tan et al., Autophagy activator AA-20 improves proteostasis and extends Caenorhabditis elegans lifespan. Proc Natl Acad Sci U S A 122, e2423455122 (2025).

16. M. K. Shimabukuro et al., Lipid-laden cells differentially distributed in the aging brain are functionally active and correspond to distinct phenotypes. Sci Rep 6, 23795 (2016).

17. Z. Q. Lan, Z. Y. Ge, S. K. Lv, B. Zhao, C. X. Li, The regulatory role of lipophagy in central nervous system diseases. Cell Death Discov 9, 229 (2023).

18. B. C. Farmer, A. E. Walsh, J. C. Kluemper, L. A. Johnson, Lipid Droplets in Neurodegenerative Disorders. Front Neurosci 14, 742 (2020).

19. L. Li et al., Visualizing Dynamic Performance of Lipid Droplets in a Parkinson’s Disease Model via a Smart Photostable Aggregation-Induced Emission Probe. iScience 21, 261– 272 (2019).

20. V. Girard et al., Abnormal accumulation of lipid droplets in neurons induces the conversion of alpha-Synuclein to proteolytic resistant forms in a Drosophila model of Parkinson’s disease. PLoS Genet 17, e1009921 (2021).

21. O. R. Brekk, J. R. Honey, S. Lee, P. J. Hallett, O. Isacson, Cell type-specific lipid storage changes in Parkinson’s disease patient brains are recapitulated by experimental glycolipid disturbance. Proc Natl Acad Sci U S A 117, 27646–27654 (2020).

22. T. Russo, M. Riessland, Lipid accumulation drives cellular senescence in dopaminergic neurons. Aging (Albany NY) 16, 11128–11133 (2024).

23. T. Vogiatzi, M. Xilouri, K. Vekrellis, L. Stefanis, Wild type alpha-synuclein is degraded by chaperone-mediated autophagy and macroautophagy in neuronal cells. J Biol Chem 283, 23542–23556 (2008).

24. S. H. Shahmoradian et al., Lewy pathology in Parkinson’s disease consists of crowded organelles and lipid membranes. Nat Neurosci 22, 1099–1109 (2019).

25. N. W. McKee et al., Lipophagy and lipid droplets in neurodegeneration. Trends Pharmacol Sci 47, 1116–1135 (2026).

26. K. Shah, D. K. Lahiri, Cdk5 activity in the brain - multiple paths of regulation. J Cell Sci 127, 2391–2400 (2014).

27. C. Ao, C. Li, J. Chen, J. Tan, L. Zeng, The role of Cdk5 in neurological disorders. Front Cell Neurosci 16, 951202 (2022).

28. Z. Tian, B. Feng, X. Q. Wang, J. Tian, Focusing on cyclin-dependent kinases 5: A potential target for neurological disorders. Front Mol Neurosci 15, 1030639 (2022).

29. A. B. Allnutt, A. K. Waters, S. Kesari, V. M. Yenugonda, Physiological and Pathological Roles of Cdk5: Potential Directions for Therapeutic Targeting in Neurodegenerative Disease. ACS Chem Neurosci 11, 1218–1230 (2020).

30. S. Singh, U. Kumar, Brain region-specific dopamine receptor changes and astrocyte activation influence tau pathology through CDK5 in Alzheimer’s disease models. J Alzheimers Dis 108, 1696–1716 (2025).

31. H. I. Mack, B. Zheng, J. M. Asara, S. M. Thomas, AMPK-dependent phosphorylation of ULK1 regulates ATG9 localization. Autophagy 8, 1197–1214 (2012).

32. N. Nandi, L. K. Tyra, D. Stenesen, H. Kramer, Acinus integrates AKT1 and subapoptotic caspase activities to regulate basal autophagy. J Cell Biol 207, 253–268 (2014).

33. N. Nandi, L. K. Tyra, D. Stenesen, H. Kramer, Stress-induced Cdk5 activity enhances cytoprotective basal autophagy in Drosophila melanogaster by phosphorylating acinus at serine(437). Elife 6, (2017).

34. T. Furuya et al., Negative regulation of Vps34 by Cdk mediated phosphorylation. Mol Cell 38, 500–511 (2010).

35. A. L. Mahul-Mellier et al., The process of Lewy body formation, rather than simply alpha-synuclein fibrillization, is one of the major drivers of neurodegeneration. Proc Natl Acad Sci U S A 117, 4971–4982 (2020).

36. B. C. Nelson et al., Lipid Droplet Binding and Oligomerization Properties of the Parkinson’s Disease Protein α-Synuclein*. Journal of Biological Chemistry 277, 6344– 6352 (2002).

37. C. Galvagnion et al., Lipid vesicles trigger α-synuclein aggregation by stimulating primary nucleation. Nature Chemical Biology 11, 229–234 (2015).

38. J. Alarcon-Gil et al., Neuroprotective and Anti-Inflammatory Effects of Linoleic Acid in Models of Parkinson’s Disease: The Implication of Lipid Droplets and Lipophagy. Cells 11, (2022).

39. J. Lim, Y. Bang, K. M. Kim, H. J. Choi, Differentiated HT22 cells as a novel model for in vitro screening of serotonin reuptake inhibitors. Front Pharmacol 13, 1062650 (2022).

40. T. Kaizuka et al., An Autophagic Flux Probe that Releases an Internal Control. Mol Cell 64, 835–849 (2016).

41. M. Mauthe et al., Chloroquine inhibits autophagic flux by decreasing autophagosome-lysosome fusion. Autophagy 14, 1435–1455 (2018).

42. E. L. Eskelinen, To be or not to be? Examples of incorrect identification of autophagic compartments in conventional transmission electron microscopy of mammalian cells. Autophagy 4, 257–260 (2008).

43. E. L. Eskelinen, A. L. Kovacs, Double membranes vs. lipid bilayers, and their significance for correct identification of macroautophagic structures. Autophagy 7, 931– 932 (2011).

44. H. T. Sakurai et al., Development of small fluorescent probes for the analysis of autophagy kinetics. iScience 26, 107218 (2023).

45. M. Ouimet et al., Autophagy regulates cholesterol efflux from macrophage foam cells via lysosomal acid lipase. Cell Metab 13, 655–667 (2011).

46. R. Singh, A. M. Cuervo, Lipophagy: connecting autophagy and lipid metabolism. Int J Cell Biol 2012, 282041 (2012).

47. D. B. Castellanos, C. A. Martin-Jimenez, F. Rojas-Rodriguez, G. E. Barreto, J. Gonzalez, Brain lipidomics as a rising field in neurodegenerative contexts: Perspectives with Machine Learning approaches. Front Neuroendocrinol 61, 100899 (2021).

48. S. Akyol et al., Lipid Profiling of Alzheimer’s Disease Brain Highlights Enrichment in Glycerol(phospho)lipid, and Sphingolipid Metabolism. Cells 10, 2591 (2021).

49. K. Huynh et al., Concordant peripheral lipidome signatures in two large clinical studies of Alzheimer’s disease. Nat Commun 11, 5698 (2020).

50. D. Paquet et al., Efficient introduction of specific homozygous and heterozygous mutations using CRISPR/Cas9. Nature 533, 125–129 (2016).

51. F. Coppede, L. Migliore, DNA damage and repair in Alzheimer’s disease. Curr Alzheimer Res 6, 36–47 (2009).

52. S. R. Labra et al., Autophagy Activators Normalize Aberrant Tau Proteostasis and Rescue Synapses in Human Familial Alzheimer’s Disease iPSC-Derived Cortical Organoids. Adv Sci (Weinh) 13, e14783 (2026).

53. M. A. Lancaster et al., Cerebral organoids model human brain development and microcephaly. Nature 501, 373–379 (2013).

54. N. T. Seyfried et al., A Multi-network Approach Identifies Protein-Specific Co-expression in Asymptomatic and Symptomatic Alzheimer’s Disease. Cell Syst 4, 60–72 e64 (2017).

55. S. Zahid, M. Oellerich, A. R. Asif, N. Ahmed, Phosphoproteome profiling of substantia nigra and cortex regions of Alzheimer’s disease patients. J Neurochem 121, 954–963 (2012).

56. K. Liu, M. J. Czaja, Regulation of lipid stores and metabolism by lipophagy. Cell Death & Differentiation 20, 3–11 (2013).

57. P. P. Hsu et al., The mTOR-regulated phosphoproteome reveals a mechanism of mTORC1-mediated inhibition of growth factor signaling. Science 332, 1317–1322 (2011).

58. M. Kanehisa, Toward understanding the origin and evolution of cellular organisms. Protein Sci 28, 1947–1951 (2019).

59. M. Kanehisa, M. Furumichi, Y. Sato, M. Kawashima, M. Ishiguro-Watanabe, KEGG for taxonomy-based analysis of pathways and genomes. Nucleic Acids Res 51, D587–D592 (2023).

60. M. Kanehisa, S. Goto, KEGG: kyoto encyclopedia of genes and genomes. Nucleic Acids Res 28, 27–30 (2000).

61. X. Wan, B. Harkavy, N. Shen, P. Grohar, L. J. Helman, Rapamycin induces feedback activation of Akt signaling through an IGF-1R-dependent mechanism. Oncogene 26, 1932–1940 (2007).

62. H. Hochegger, S. Takeda, T. Hunt, Cyclin-dependent kinases and cell-cycle transitions: does one fit all? Nat Rev Mol Cell Biol 9, 910–916 (2008).

63. K. Zheng, Z. He, K. Kitazato, Y. Wang, Selective Autophagy Regulates Cell Cycle in Cancer Therapy. Theranostics 9, 104–125 (2019).

64. C. Wang et al., Phosphorylation of ULK1 affects autophagosome fusion and links chaperone-mediated autophagy to macroautophagy. Nat Commun 9, 3492 (2018).

65. Z. H. Cheung, N. Y. Ip, Cdk5: a multifaceted kinase in neurodegenerative diseases. Trends Cell Biol 22, 169–175 (2012).

66. H. L. De Bondt et al., Crystal structure of cyclin-dependent kinase 2. Nature 363, 595–602 (1993).

67. C. Tarricone et al., Structure and regulation of the CDK5-p25(nck5a) complex. Mol Cell 8, 657–669 (2001).

68. R. L. Rich, D. G. Myszka, Advances in surface plasmon resonance biosensor analysis. Curr Opin Biotechnol 11, 54–61 (2000).

69. M. Mapelli et al., Mechanism of CDK5/p25 binding by CDK inhibitors. J Med Chem 48, 671–679 (2005).

70. O. Trott, A. J. Olson, AutoDock Vina: improving the speed and accuracy of docking with a new scoring function, efficient optimization, and multithreading. J Comput Chem 31, 455–461 (2010).

71. K. Dong, X. Wang, X. Yang, X. Zhu, Binding mechanism of CDK5 with roscovitine derivatives based on molecular dynamics simulations and MM/PBSA methods. J Mol Graph Model 68, 57–67 (2016).

72. P. D. Jeffrey et al., Mechanism of CDK activation revealed by the structure of a cyclinA-CDK2 complex. Nature 376, 313–320 (1995).

73. J. Lee et al., Acetylation of cyclin-dependent kinase 5 is mediated by GCN5. Biochem Biophys Res Commun 447, 121–127 (2014).

74. W. Filgueira de Azevedo, Jr., R. T. Gaspar, F. Canduri, J. C. Camera, Jr., N. J. Freitas da Silveira, Molecular model of cyclin-dependent kinase 5 complexed with roscovitine. Biochem Biophys Res Commun 297, 1154–1158 (2002).

75. M. Otyepka, I. Bartova, Z. Kriz, J. Koca, Different mechanisms of CDK5 and CDK2 activation as revealed by CDK5/p25 and CDK2/cyclin A dynamics. J Biol Chem 281, 7271–7281 (2006).

76. C. R. Coxon et al., Cyclin-Dependent Kinase (CDK) Inhibitors: Structure-Activity Relationships and Insights into the CDK-2 Selectivity of 6-Substituted 2-Arylaminopurines. J Med Chem 60, 1746–1767 (2017).

77. J. S. Ahn et al., Defining Cdk5 ligand chemical space with small molecule inhibitors of tau phosphorylation. Chem Biol 12, 811–823 (2005).

78. S. L. Rath, S. Senapati, Molecular basis of differential selectivity of cyclobutyl-substituted imidazole inhibitors against CDKs: insights for rational drug design. PLoS One 8, e73836 (2013).

79. L. Meijer et al., Biochemical and cellular effects of roscovitine, a potent and selective inhibitor of the cyclin-dependent kinases cdc2, cdk2 and cdk5. Eur J Biochem 243, 527–536 (1997).

80. P.-C. Pao, L.-H. Tsai, Three decades of Cdk5. Journal of Biomedical Science 28, 79 (2021).

81. C. P. Poore et al., Cdk5-mediated phosphorylation of delta-catenin regulates its localization and GluR2-mediated synaptic activity. J Neurosci 30, 8457–8467 (2010).

82. T. B. Shea et al., Cdk5 regulates axonal transport and phosphorylation of neurofilaments in cultured neurons. J Cell Sci 117, 933–941 (2004).

83. S. Yukio et al., Fyn and Cdk5 Mediate Semaphorin-3A Signaling, Which Is Involved in Regulation of Dendrite Orientation in Cerebral Cortex. Neuron 35, 907–920 (2002).

84. L. A. Volpicelli-Daley et al., Exogenous alpha-synuclein fibrils induce Lewy body pathology leading to synaptic dysfunction and neuron death. Neuron 72, 57–71 (2011).

85. H. Fujiwara et al., alpha-Synuclein is phosphorylated in synucleinopathy lesions. Nat Cell Biol 4, 160–164 (2002).

86. J. L. Webb, B. Ravikumar, J. Atkins, J. N. Skepper, D. C. Rubinsztein, Alpha-Synuclein is degraded by both autophagy and the proteasome. J Biol Chem 278, 25009–25013 (2003).

87. S. K. Mak, A. L. McCormack, A. B. Manning-Bog, A. M. Cuervo, D. A. Di Monte, Lysosomal degradation of alpha-synuclein in vivo. J Biol Chem 285, 13621–13629 (2010).

88. E. Sidransky et al., Multicenter analysis of glucocerebrosidase mutations in Parkinson’s disease. N Engl J Med 361, 1651–1661 (2009).

89. N. Malik et al., Induction of lysosomal and mitochondrial biogenesis by AMPK phosphorylation of FNIP1. Science 380, eabj5559 (2023).

90. S. Yang et al., Regulation of TFEB nuclear localization by HSP90AA1 promotes autophagy and longevity. Autophagy 19, 822–838 (2023).

91. S. T. Kumar, S. Donzelli, A. Chiki, M. M. K. Syed, H. A. Lashuel, A simple, versatile and robust centrifugation-based filtration protocol for the isolation and quantification of alpha-synuclein monomers, oligomers and fibrils: Towards improving experimental reproducibility in alpha-synuclein research. J Neurochem 153, 103–119 (2020).

92. A.-L. Mahul-Mellier et al., The making of a Lewy body: the role of alpha-synuclein post-fibrillization modifications in regulating the formation and the maturation of pathological inclusions. bioRxiv, 500058 (2018).

93. H. Schägger, Tricine-SDS-PAGE. Nature protocols 1, 16–22 (2006).

94. R. Grosse-Kunstleve, N. Sauter, N. Moriarty, P. Adams, The Computational Crystallography Toolbox: Crystallographic algorithms in a reusable software framework. Computational Crystallography Toolbox J. Appl. Cryst. 126±136 Journal 35, 126–136 (2002).

95. P. Eastman et al., OpenMM 8: Molecular Dynamics Simulation with Machine Learning Potentials. The Journal of Physical Chemistry B 128, 109–116 (2024).

96. R. T. McGibbon et al., MDTraj: A Modern Open Library for the Analysis of Molecular Dynamics Trajectories. Biophys J 109, 1528–1532 (2015).

97. R. J. Gowers et al., in SciPy. (2016).

98. N. Michaud-Agrawal, E. J. Denning, T. B. Woolf, O. Beckstein, MDAnalysis: a toolkit for the analysis of molecular dynamics simulations. J Comput Chem 32, 2319–2327 (2011).

99. S. Zhang et al., ProDy 2.0: increased scale and scope after 10 years of protein dynamics modelling with Python. Bioinformatics (Oxford, England) 37, 3657–3659 (2021).

100. P. Virtanen et al., SciPy 1.0: fundamental algorithms for scientific computing in Python. Nature Methods 17, 261–272 (2020).

101. C. R. Harris et al., Array programming with NumPy. Nature 585, 357–362 (2020).

102. W. McKinney, in SciPy. (2010).

103. J. D. Hunter, Matplotlib: A 2D Graphics Environment. Computing in Science & Engineering 9, 90–95 (2007).

104. L. L. C. Schrodinger. (2010).

105. S. Ghatak et al., NitroSynapsin ameliorates hypersynchronous neural network activity in Alzheimer hiPSC models. Mol Psychiatry 26, 5751–5765 (2021).

106. S. Ghatak et al., Mechanisms of hyperexcitability in Alzheimer’s disease hiPSC-derived neurons and cerebral organoids vs isogenic controls. Elife 8, (2019).

107. S.-J. Yoon et al., Reliability of human cortical organoid generation. Nature Methods 16, 75–78 (2019).

108. Q. Luo et al., Targetable leukaemia dependency on noncanonical PI3Kγ signalling. Nature 630, 198–205 (2024).

109. J. Navarrete-Perea, Q. Yu, S. P. Gygi, J. A. Paulo, Streamlined Tandem Mass Tag (SL-TMT) Protocol: An Efficient Strategy for Quantitative (Phospho)proteome Profiling Using Tandem Mass Tag-Synchronous Precursor Selection-MS3. J Proteome Res 17, 2226– 2236 (2018).

110. J. Rappsilber, Y. Ishihama, M. Mann, Stop and go extraction tips for matrix-assisted laser desorption/ionization, nanoelectrospray, and LC/MS sample pretreatment in proteomics. Anal Chem 75, 663–670 (2003).

111. D. K. Schweppe et al., Characterization and Optimization of Multiplexed Quantitative Analyses Using High-Field Asymmetric-Waveform Ion Mobility Mass Spectrometry. Anal Chem 91, 4010–4016 (2019).

112. D. K. Schweppe, S. F. Rusin, S. P. Gygi, J. A. Paulo, Optimized Workflow for Multiplexed Phosphorylation Analysis of TMT-Labeled Peptides Using High-Field Asymmetric Waveform Ion Mobility Spectrometry. J Proteome Res 19, 554–560 (2020).

113. D. Kessner, M. Chambers, R. Burke, D. Agus, P. Mallick, ProteoWizard: open source software for rapid proteomics tools development. Bioinformatics 24, 2534–2536 (2008).

114. A. T. Kong, F. V. Leprevost, D. M. Avtonomov, D. Mellacheruvu, A. I. Nesvizhskii, MSFragger: ultrafast and comprehensive peptide identification in mass spectrometry-based proteomics. Nat Methods 14, 513–520 (2017).

115. M. The, M. J. MacCoss, W. S. Noble, L. Käll, Fast and Accurate Protein False Discovery Rates on Large-Scale Proteomics Data Sets with Percolator 3.0. J Am Soc Mass Spectrom 27, 1719–1727 (2016).

116. A. I. Nesvizhskii, A. Keller, E. Kolker, R. Aebersold, A statistical model for identifying proteins by tandem mass spectrometry. Anal Chem 75, 4646–4658 (2003).

117. H.-Y. Chang et al., Analysis of isobaric quantitative proteomic data using TMT-Integrator and FragPipe computational platform. bioRxiv, 2025.2005.2027.656447 (2025).

118. M. E. Ritchie et al., limma powers differential expression analyses for RNA-sequencing and microarray studies. Nucleic Acids Research 43, e47–e47 (2015).

119. K. Engholm-Keller et al., The temporal profile of activity-dependent presynaptic phospho-signalling reveals long-lasting patterns of poststimulus regulation. PLoS Biol 17, e3000170 (2019).

120. P. V. Hornbeck et al., 15 years of PhosphoSitePlus®: integrating post-translationally modified sites, disease variants and isoforms. Nucleic Acids Res 47, D433–d441 (2019).

121. Y. Perez-Riverol et al., The PRIDE database at 20 years: 2025 update. Nucleic Acids Res 53, D543–D553 (2025).

